# Iterative deep learning-design of human enhancers exploits condensed sequence grammar to achieve cell type-specificity

**DOI:** 10.1101/2024.06.14.599076

**Authors:** Christopher Yin, Sebastian Castillo Hair, Gun Woo Byeon, Peter Bromley, Wouter Meuleman, Georg Seelig

## Abstract

An important and largely unsolved problem in synthetic biology is how to target gene expression to specific cell types. Here, we apply iterative deep learning to design synthetic enhancers with strong differential activity between two human cell lines. We initially train models on published datasets of enhancer activity and chromatin accessibility and use them to guide the design of synthetic enhancers that maximize predicted specificity. We experimentally validate these sequences, use the measurements to re-optimize the predictor, and design a second generation of enhancers with improved specificity. Our design methods embed relevant transcription factor binding site (TFBS) motifs with higher frequencies than comparable endogenous enhancers while using a more selective motif vocabulary, and we show that enhancer activity is correlated with transcription factor expression at the single cell level. Finally, we characterize causal features of top enhancers via perturbation experiments and show enhancers as short as 50bp can maintain specificity.

## Introduction

Enhancers are a class of *cis*-regulatory elements (CREs) that exist in the non-coding regions of the human genome and significantly determine differential gene expression across cell types. Transcription Factor Binding Sites (TFBSs), short sequence fragments recognized and bound by corresponding transcription factors (TFs), are the core functional units within an enhancer sequence^1^ but occupy only a small fraction of the enhancer footprint, which typically spans hundreds or even thousands of base pairs^2^. Overall enhancer activity is controlled by a cell type-specific and incompletely understood *cis*-regulatory code encompassing TFBS identity, abundance, relative positioning, and flanking sequences, among other grammatical elements^1^.

One major obstacle in deciphering this code is the difficulty of identifying functional enhancers and associating them with the potentially distal gene(s) they regulate. Chromatin accessibility has shown to be a valuable indicator of regulatory elements^3–5^, and genome-scale sequencing workflows mapping DNase I Hypersensitive Sites (DHSs) are providing increasingly detailed tissue and cell type-specific atlases of these elements^6–8^. However, although these data form a very important part of the puzzle, our understanding of the mapping from chromatin state to enhancer activity remains incomplete; not every element carrying the requisite hallmarks appears to encode an active enhancer and drive cell type-specific gene expression^2^.

Enhancers that impact gene expression in a specific cell type are powerful tools for basic biology and can increase the specificity of gene therapies, thereby reducing side effects. Chromatin accessibility data are a common starting point for curating putative cell type-specific enhancers due to their wide availability, although they often do not provide direct evidence of expression effects. Alternatively, enhancers can be selected from Massively Parallel Reporter Assays (MPRAs)^2^ which provide a direct readout of the ability of a DNA fragment to drive gene expression of a minimal promoter in the context of a plasmid. However, in spite of recent progress in lentiviral delivery^9^ and single cell readout^10,11^ of MPRAs, the assessment of large reporter libraries in complex tissues remains a bottleneck, and MPRA data are only available for a limited number of cell types. Moreover, an MPRA is not guaranteed to identify an enhancer with the desired properties, as the range of outcomes is constrained by the number and predetermined identities of the tested sequences.

Current state-of-the-art approaches to confront the limitation of purely experimental screens involve deep learning models trained on large-scale genomic and/or MPRA datasets. Such models are theoretically capable of learning a sequence-to-function mapping that captures underlying biological principles^12–17^, and can thereby guide the design of synthetic enhancers with targeted activity levels. DaSilva *et al.* and Lal *et al.* demonstrated synthetic cell type-specific enhancer design *in silico*^18,19^. De Almeida *et al.* experimentally validated synthetic enhancer designs, initially using STARR-seq to prove enhancer activity in a single *Drosophila* developmental cell type^15^, then targeting enhancers to four distinct tissue types in the *Drosophila* embryo and confirming specificity with *in vivo* assays^20^. In these studies, sequences were randomly generated and those with the highest model-predicted activity and/or specificity were selected for testing. Taskiran *et al.* also showed enhancer design in *Drosophila* cell types but used model-directed mutagenesis and Generative Adversarial Networks (GANs); they also demonstrated cell type-specific enhancer design in melanoma-derived human cell lines^21^ and validated a small subset of sequences via fluorescence-based assays. Gosai *et al.* scaled synthetic enhancer design and validation using MPRA-trained models in conjunction with generative methods to design a library of *de novo* enhancers for specific activity in a set of three cell lines (HepG2, K562, SK-N-SH)^22^.

In this work we advance the nascent field of deep learning-based enhancer design through validation and extensive characterization of a high-throughput, multimodal, and iterative design process. We train models on two different types of previously published data: reporter-based (MPRA) and accessibility-based (DHS). We then apply several distinct strategies to design libraries of candidate enhancers that maximize differential enhancer activity. We test these synthetic enhancers in human cell lines (HepG2 and K562), then retrain our model with these data and repeat the design-build-test cycle to show that iterative retraining results in dramatic performance improvements. Strikingly, it does so even with a relatively small amount of data. Such an iterative “small data” approach may be more readily adapted to selecting enhancers for specific cell types *in vivo*, rather than an approach that requires testing hundreds of thousands of sequences for model training. Model interpretation and motif analysis techniques reveal how the information content encoded in the enhancers evolves with each round of model training towards a more selective TFBS vocabulary and higher density syntax. Additionally, we perform motif ablations for a subset of enhancers to experimentally and computationally quantify the *cis*- regulatory code. Finally, we perform a single cell MPRA (scMPRA) to quantify cell-to-cell variation in enhancer activity and show correlated activity with cognate TF expression levels.

## Results

### Deep learning predictors enable design of cell type-specific enhancers from MPRA data

Enhancer MPRAs measure the ability of a DNA fragment to drive reporter gene expression and thus provide a functional readout of the regulatory activity of a putative enhancer. Here, we set out to capture sequence-encoded determinants of cell type-specific enhancer activity, and thereby guide the design of synthetic enhancers with tailor-made properties. To this end, we trained neural network predictors on the previously published Sharpr-MPRA dataset^23^. This dataset consists of two replicates of enhancer strength measurements (log_2_(mRNA counts / DNA counts)) from 467,000 145 nt-long candidate enhancers extracted from accessible genomic regions (**Fig. 1A**), cloned into reporter plasmids upstream of a minimal promoter (minP), and assayed in HepG2 and K562 cells. We refer to enhancer strength in each cell line as log2FC_HepG2_ and log2FC_K562_, and define the differential strength or specificity log2FC_H2K_ as log2FC_HepG2_ - log2FC_K562_. We filtered out sequences with low read coverage and low specificity, and retained 29,891 sequences for model training (**Fig. S1A, Methods**). We refer to this initial dataset as R0-MPRA (**Fig. 1B**).

**Fig. 1.**
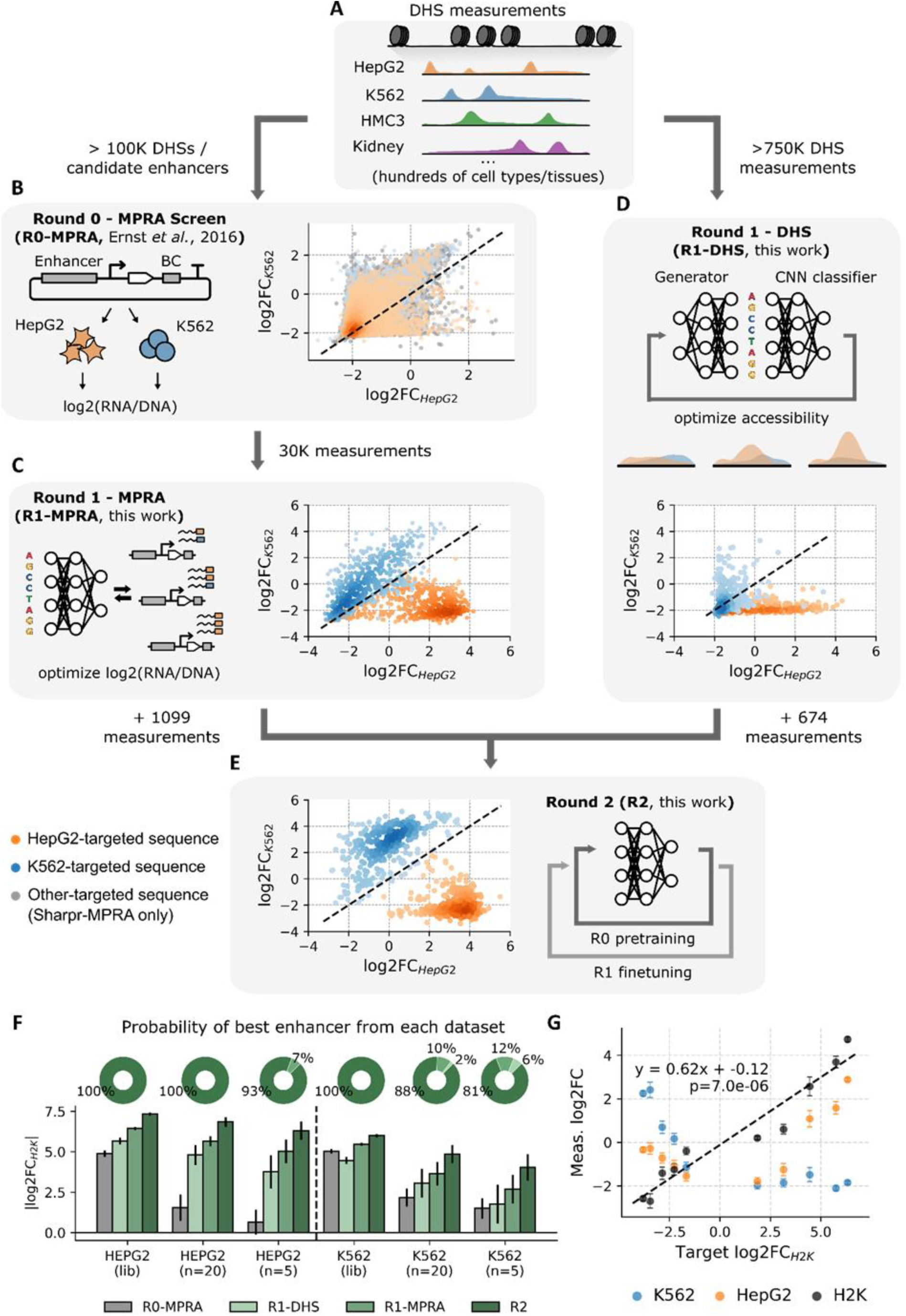
Multi-round model-based enhancer design results in cell type-specific sequences with improved performance. **A-E,** Overview of iterative design workflow. Accessibility measurements are first used to identify putative regulatory elements in the genome (**A**). The Sharpr-MPRA^23^ assayed >100k of such candidates in HepG2 and K562 via MPRA, which we filtered for read depth and enhancer strength (R0-MPRA, **B**). We designed and tested sequences using deep learning models trained on these MPRA data (R1-MPRA, **C**) or directly on accessibility (R1-DHS, **D**), and used the results to improve our models and further increase performance (R2, **E**). Individual markers correspond to measured enhancer strength in HepG2 (x-axis) vs K562 (y-axis). Dashed line: y = x. For Sharpr-MPRA sequences in **B**, target cell type is matched to the cell type of the accessibility dataset from which it was derived; cell types other than HepG2 and K562 are colored gray. **F,** Strength and origin of the most specific enhancers across design rounds, as a function of library size. Top row, bootstrap-estimated probability that the most specific enhancer across all tested libraries will come from a given design round, when each entire library is considered (lib), or with simulated sizes of n=20 and n=5. Bottom row: |log2FC_H2K_| of most specific enhancer in each design round, library size, and target cell type. For “lib” columns, bar height and error bars are the mean and DEseq2-estimated standard error of individual measurements; for “n=20” and “n=5” bar height and error bars are the mean and standard deviation across 10,000 bootstrap simulations. **G**, Performance of enhancers designed for intermediate target specificities. Markers and error bars are the mean and standard deviation across 20 (intermediate targets) or 110 (largest HepG2 or K562 target values) sequences designed for a given target value. The line of best fit is plotted (dashed black).

To avoid overreliance on any single model or design approach, we trained a total of 120 multitask convolutional neural network (CNN) models to predict log2FC_HepG2_ and log2FC_K562_ from one hot-encoded DNA sequence input (**Methods, Fig. S1B**), exploring several variations of model training all achieving prediction-measurement correlation on par with the inter-replicate correlation of R0-MPRA (**Methods, Fig. S1C)**. We then applied three different design methods (Simulated Annealing, Fast SeqProp^24^, Deep Exploration Networks^25^, **Methods**) to generate 1037 *de novo* candidate enhancers that maximize predicted differential enhancer activity. We verified that these sequences were highly dissimilar to one another and to the Sharpr-MPRA dataset (**Fig. S1D**). As controls, we re-synthesized 100 sequences from Sharpr-MPRA selected to span the entire range of activities. The reverse complements of these 100 control sequences were synthesized as well to test the impact of enhancer orientation. Collectively, these MPRA-guided designs and associated controls are referred to as R1-MPRA and the detailed library composition is shown in **Table S1**.

We experimentally quantified the strength of our synthetic enhancers and control sequences in the R1-MPRA library using the reporter construct and cell lines from Sharpr-MPRA^23^ (**Methods**). Our measurements were highly consistent across replicates (**Fig. S1E**), and Sharpr controls and their reverse complements were well correlated with measurements in the original study (**Fig. S1F,G**). mRNA expression from designed enhancers overwhelmingly matched their target specificity: most sequences designed to target HepG2 resulted in higher expression in HepG2 compared to K562 (median log2FC_H2K_ = 3.28, 9.7x higher HepG2 expression), and vice versa (median log2FC_H2K_ = −1.22, 2.3x higher K562 expression). Furthermore, many sequences achieved cell type-specificity not merely via weak activity in the off-target cell line, but via strong activity in the target cell line (**Fig. 1C**). In general, HepG2-targeted designs exhibited higher differential strength, which may reflect the lower replicate correlation observed in the K562 measurements of the training data (**Fig. S1A**). Similarly, 113/523 (22%) synthetic enhancers showed greater log2FC_H2K_ than the most HepG2-specific control (4.80), but only 2/514 (0.004%) synthetic sequences surpassed the most K562-specific control sequence (−5.03).

### Deep learning predictors enable design of cell type-specific enhancers from chromatin accessibility

Unlike enhancer MPRA data, DNA accessibility measurements are available for a plethora of cell and tissue types, providing a promising starting point for the selection and design of enhancer elements for biological contexts where MPRA data are not available. We wanted to assess to what extent models trained on accessibility data alone can be used to generate functional enhancers that exhibit strong and specific activity in human cells.

To this end, we trained a Generative Adversarial Network (GAN)^26^ model using a compendium of 750k+ endogenous example elements obtained from large-scale chromatin accessibility assays encompassing >400 distinct cell and tissue types^8^. Briefly, in these assays chromatin is digested using the DNase I endonuclease, preferentially cleaving sites that are accessible to DNA binding factors and thus allowing the genome-wide identification of DNase I Hypersensitive Sites (DHSs). After training, the GAN produced *de novo* sequences resembling endogenous genomic accessible elements (**Fig. S2A,B**, **Methods**), but not explicitly geared towards specific cellular contexts. To maximize cell type-specificity, we subsequently tuned generated sequences using a separate classifier model trained on a subset of sequences specifically accessible in HepG2, K562, or otherwise (library referred to as R0-DHS, **Methods**). The tuning process is relatively stable, largely retaining the original sequence identity of generated sequences across many tuning iterations (**Fig. S2C,D**, **Methods**); and has limited impact on sequence characteristics (**Fig. S2E,F**). Nonetheless, during tuning we observe a steady increase of the number of sequences containing TF motifs relevant^8^ for our cellular contexts of interest (**Fig. S2G,H**). After this tuning process, for each cell type we selected 300 designed sequences plus 37 positive-control genomic sequences for subsequent experimental validation; we refer to this library as R1-DHS (**Methods**).

Strikingly, despite being designed for chromatin accessibility only, R1-DHS sequences strongly drove cell type-specific gene expression (**Fig. 1D**). As in the MPRA-based approach, HepG2 designs were more successful, with 46/300 (15%) designs exceeding the specificity of the best R0-DHS control (3.75), compared to 9/300 (3%) designs exceeding the specificity of the best K562 control (−3.45). The median log2FC_H2K_ of R1-DHS sequences was 1.71 (3.3x higher HepG2 expression) for HepG2 designs, and 0.04 for K562 designs.

R1-DHS sequences on average exhibited lower activity than R1-MPRA sequences (**Fig. 1C,D,F, Fig. S4A**), consistent with R1-MPRA designs being generated from models trained directly on enhancer activity data. In HepG2-targeted designs median log2FC_H2K_ was 1.71 in R1-DHS vs 3.28 in R1-MPRA (p=2.8e-16, Wilcoxon rank sum); and in K562-targeted designs median log2FC_H2K_ was 0.04 vs −1.22 (p=2.5e-34, Wilcoxon rank sum). We note that a direct comparison between R1-DHS and R1-MPRA may not be meaningful because of differences in the data and design approaches. For example, the accessibility-based designs explicitly aim to minimize accessibility in cell types other than the two target cell lines; no such constraint was used in the MPRA designs, because activity measurements were only available for the two targets. Moreover, accessibility-based designs are regularized to be similar to native accessible elements while the MPRA-based designs do not use such regularization and may in fact encourage the generation of sequences that look more extreme than those encountered in the training dataset.

### Iterative retraining and design results in improved enhancer specificity and precision

Next, we asked whether design performance could be improved by retraining models with R1-MPRA and R1-DHS data. This was motivated by the observation that despite having high predicted specific activity or accessibility, not all designed sequences in the R1 libraries were measured to be strongly specific, suggesting the presence of regulatory grammar that the initial models do not capture. Additionally, compared to the original datasets, the R1 libraries contain a much higher fraction of positive (i.e. highly active) examples and thus have the potential to improve model performance in the most relevant part of the design space. Practically, while it remains challenging or even impossible to perform MPRAs in specific primary cell types at scale, testing a few hundred or even a few thousand sequences in complex tissues in a single cell MPRA (scMPRA) format is within reach^10,11^. Thus it is important to understand if retraining with a relatively limited dataset, but one that is highly enriched for functional sequences, can iteratively increase the success rate of enhancer designs.

We evaluated training models with R1 measurements using two distinct strategies. First, we sought to improve performance of a CNN ensemble model by pretraining on R0-MPRA and performing additional training iterations on R1-MPRA and R1-DHS (“M0+1” models, **Methods**). Second, we trained an identical CNN ensemble on both R1 datasets from scratch without R0-MPRA pretraining (“M1” models), to test whether a small dataset enriched in functional sequences can be sufficient for model-based design. Surprisingly, M1 models nearly matched M0+1 prediction performance on a held-out R1-MPRA test set (**Fig. S3A**). We then generated a new set of 690 sequences to maximize target specificity using both the M0+1 and M1 ensembles (**Methods**). Library diversity was confirmed as above (**Fig. S3B**). As controls, we also included the top 5 enhancers in each cell line from R1-MPRA, as well as 200 randomly sampled R1-MPRA sequences. We refer to this set of designed and control sequences as R2 (**Table S2**).

We experimentally assayed R2 enhancers as before and with equivalently strong replicate correlation (**Fig. S3C**), observing dramatically higher median specificities of both HepG2-targeted and K562-targeted sequences in comparison to the R1 libraries (**Fig. S4A**). For HepG2-targeted sequences, median expression was 46.2-fold higher in HepG2 cells compared to K562 cells (log2FC_H2K_ = 5.53); and for K562-targeted sequences median expression was 6.7-fold higher in the target cell line (log2FC_H2K_ = −2.74) (**Fig. 1E**). Evaluating the most specific sequences from each round, iterative improvement was observed in both cell types. For HepG2 designs the best log2FC_H2K_ increased from 4.87 (R0) to 6.44/5.66 (R1-MPRA/R1-DHS) to 7.34 (R2), with significant improvement across all consecutive rounds. For K562 designs the best log2FC_H2K_ decreased from −5.03 to −5.47/−4.45 to −5.99, with significant improvement between R1 and R2 (**Fig. 1F, bottom**).

To simulate *in vivo* experimental constraints, where library throughput is greatly reduced compared to MPRAs^5^, we also estimated the expected best enhancer performance for smaller library sizes (n=5, 20) using bootstrap sampling of R2. Enhancer performance remained strong for R2 across all library sizes. Testing only 5 random HepG2 enhancers yielded on average a best log2FC_H2K_ of 6.31±0.57 (mean±standard deviation), exceeding all R0 enhancers (**Fig. 1F, bottom**). We conducted identical bootstrap simulations in R0 and R1 and found that for a library size of 5, R2 enhancers outperformed those in all other libraries 92.6% (HepG2) or 81.1% (K562) of the time. When randomly selecting 20 sequences, R2 enhancers outperformed the other libraries 99.4% (HepG2) or 88.0% (K562) of the time; R1 enhancers were superior in the remaining cases, whereas R0-MPRA never outperformed any of the others (**Fig. 1F, top**). Thus, even for small sample sizes, synthetic enhancers greatly outperform those from the initial training data.

Additionally, we asked whether the improvement in model performance from iterative retraining would allow us to more precisely control enhancer specificity. Such an ability to tune target expression while maintaining specificity may be desirable in contexts where e.g. overexpression of a gene leads to deleterious and difficult to predict knock-on effects^27–29^. We used M0+1 models to design sequences targeting five different levels across a range of log2FC_H2K_ values and experimentally tested them in the R2 library, observing good relative accuracy (r^2^ = 0.93) despite only designing twenty enhancers per intermediate target (**Fig. 1G**). Although measured expression levels were generally slightly lower than predicted, we observe an almost perfect monotonic relationship between target and mean measured log2FC_H2K_.

### Comparing sublibraries of synthetic enhancers suggests optimal design practices

Across all synthetic libraries we explored a range of model architectures, design algorithms, and design objectives. Here, we compare design performance under these approaches to identify best practices and develop recommendations for future work.

First, we found that model performance was positively associated with enhancer performance. Even though sets with perfectly matched model type and design method do not exist across R1-MPRA and R2, the latter were designed with more accurate models and significantly outperformed those in R1-MPRA (**Fig. S1C, Fig. S3A, Fig. 1F, Fig. S4A)**. Comparing the median specificity of the best combination of model type and design method from each round, we observe significantly better log2FC_H2K_ in R2 vs. R1-MPRA for both HepG2 (5.80 vs 4.27, p=1.8e-22; Wilcoxon rank sum) and K562 (−3.08 vs −2.08, p=8.0e-3) designs. This trend holds for model variations within each design round. Among R2 enhancers, M0+1 models achieved significantly better median specificity in both cell lines (HepG2 median = 5.80 vs 5.42, p = 7.4e-3; K562 median = −3.08 vs −1.98, p=4.4e-9) (**Fig. S4B**). Within R1-MPRA enhancers, we also found that design performance tracked model accuracy. Three model variations were explored in R1-MPRA, all implementing the same core architecture: “Single” models trained on a constant training/validation/test data split; “Boot” models trained on randomly resampled bootstraps of the merged training and validation data splits; and “Ensemble” models formed by averaging the outputs of 10 Boot models. In HepG2-targeted designs, median specificity was significantly higher with the most accurate Ensemble models (4.27) compared to Single models (3.93, p = 1.2e-2), which in turn was higher than the least accurate Boot models (1.71, p = 9.2e-5); though no significant differences were observed between model types in K562-targeted designs (**Fig. S4C**).

Second, in contrast to the above analysis we found that all sequence design methods used in R1-MPRA (DENs, Fast SeqProp, Simulated Annealing) performed comparably well. For HepG2 designs there was a significantly higher median log2FC_H2K_ in Fast SeqProp- vs DEN-generated enhancers (3.23 vs 1.71, p=4.5e-3), but otherwise no significant differences were observed between design methods (**Fig. S4D**). Taken together, these observations suggest that predictor performance was the “rate-limiting” factor in our design approach. This motivated the exclusive use of Fast SeqProp for the R2 designs: given similar performance, we chose the approach that was least computationally onerous.

Third, we initially hypothesized that enhancers designed to maximize specificity unbounded may result in unrealistic sequences with poor performance due to overfitting, but found this was not the case. In R2 we explicitly tested this by designing sequences that either maximized |log2FC_H2K_| unbounded, or clipped to a threshold (1.1X the maximum value predicted by the M0+1 model on R1-MPRA sequences, **Table S2**). Sequences designed with the unbounded objective outperformed those designed with the clipped objective in both cell types (HepG2 median specificity = 5.80 vs 4.96, p=5.6e-9; K562 median specificity = −3.08 vs −2.59, p=8.0e-5) (**Fig. S4B**). These results suggest that our models are capable of extrapolating beyond the training data.

Finally, we asked whether there is a trade-off between maximizing specificity and on-target activity. To investigate if enforcing low off-target activity caps on-target activity, we designed 20 enhancers each to maximize (“Max1”) or minimize (“Min1”) activity in only one cell type, regardless of the other (**Table S2**). Specificity was low for all Min1 designs, which had comparably low activity in both cell types (**Fig. S4E,F**). While target cell type activity was similarly high across all Max1 designs, the specificity of these sequences varied significantly (**Fig. S4E,G**). Notably, this loss in specificity was not balanced out by a gain in absolute expression levels, as the most specific R2 enhancers obtained equivalent levels of target cell type activity (**Fig. S6G-I**).

### Designed sequences evolve more compact TFBS motif grammar

To elucidate how DL-designed enhancer grammar differed from its endogenous counterpart, we scanned all sequences for matches to the JASPAR2022 Core Vertebrate database of 137 TFBS motif clusters^30,31^, discovering 67 (R0-MPRA), 48 (R0-DHS), 50 (R1-MPRA), 41 (R1-DHS), and 23 (R2) unique motif clusters in each respective library. Motif density increased significantly over successive design rounds, and was lower in DHS-based vs MPRA-based libraries, indicating GAN designs were possibly more similar to natural sequences than designs produced via the other methods (**Fig. 2A**). Across all designed libraries motif density was positively associated with specificity. In R2, HepG2-targeted enhancers showed higher baseline activity at low motif densities, suggesting specificity was achieved with a smaller number of strongly activating motifs; whereas K562-targeted enhancers generally achieved specificity with a larger number of individually weaker motifs (**Fig. 2B,C**).

**Fig. 2.**
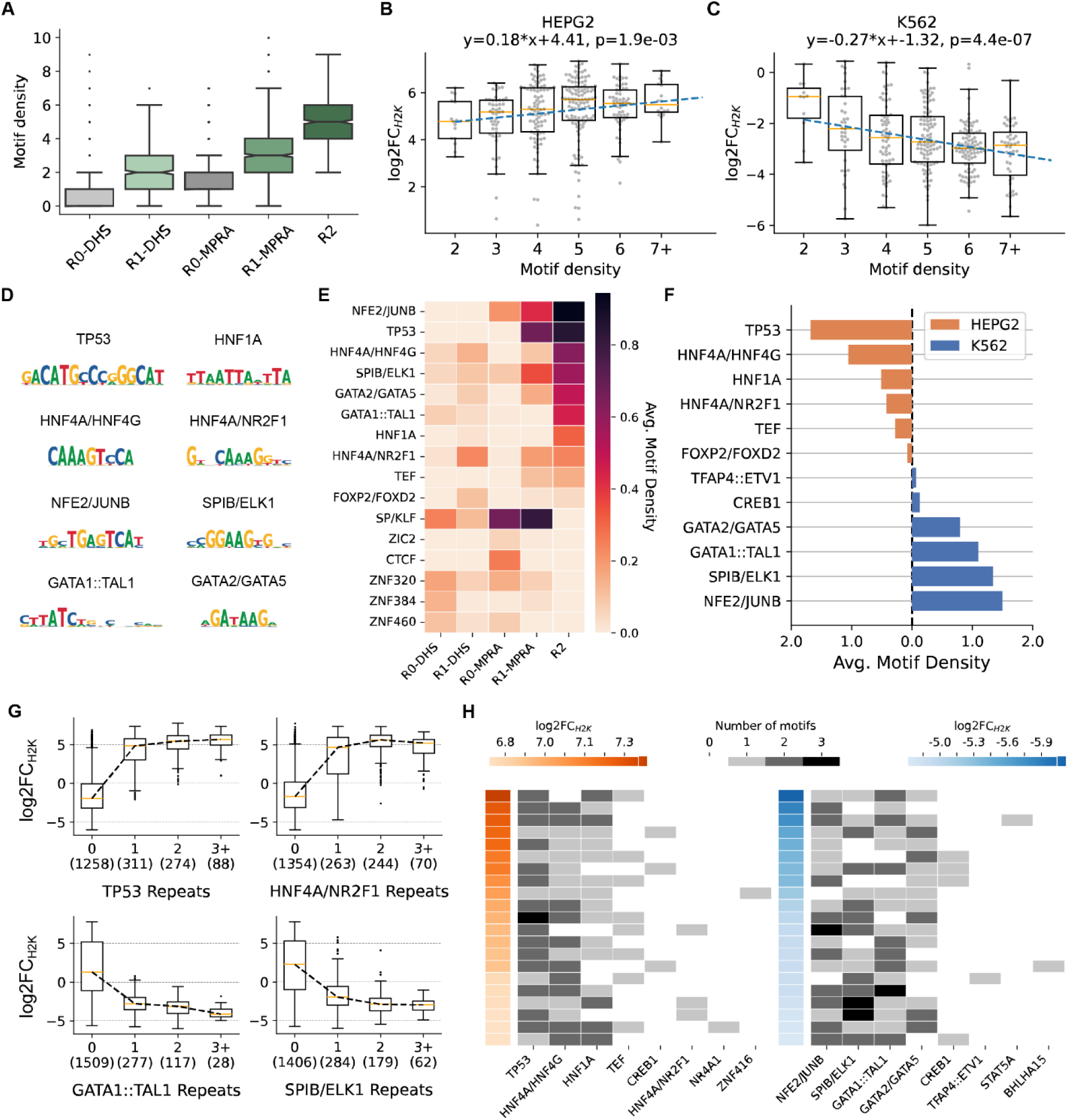
Synthetic enhancers exhibit more compressed TFBS motif grammar than natural enhancers. **A,** Motif density (number of JASPAR2022 motif cluster matches per sequence, **Methods**) of synthetic enhancers exceeds genomic motif density and increases across consecutive rounds of design. **B-C,** In R2 designs, motif density is positively correlated with enhancer specificity. Each marker corresponds to an enhancer targeted to HepG2 (**B**) or K562 (**C**). Enhancers with motif density >=7 grouped together due to low sample size. No R2 enhancers had motif density < 2. Line of best fit plotted in blue. **D,** PWMs of motifs associated with cell type-specific enhancer activity. Motifs were extracted from the most specific R1 (|log2FC_H2K_|>2) and R2 (|log2FC_H2K_|>3) sequences using STREME^43^. **E,** Average motif density (number of motif occurrences / number of sequences) within each design round for a subset of motifs with high enrichment in at least one library. **F,** Average motif density in the most cell type-specific R2 sequences (log2C_H2K_ > 3, left; log2C_H2K_ < −3, right) for the most differentially enriched motifs. **G,** Boxplots of log2FC_H2K_ measurements for R2 enhancers grouped by motif multiplicity (number of discrete instances of a given motif type in the same sequence), for a selection of motifs that show an association between multiplicity and increased specificity. Sequences with multiplicity >=3 are grouped together due to low sample sizes. For each x-value, the number of enhancers is reported in parentheses. **H**, Motif content of top 5% most-specific R2 enhancers for HepG2 designs (left), K562 designs (right). Each row represents a single enhancer sequence (sorted by descending specificity), columns indicate different motifs, cell color indicates motif multiplicity. log2FC_H2K_ plotted as a colored cell alongside corresponding sequence rows.

Motif composition of the synthetic libraries significantly evolved over successive design iterations, and was markedly different from their genomic counterparts (**Fig. 2D,E**). As a striking example, TP53 was modestly represented in the R0-MPRA dataset (41 TP53 motif hits across 29,891 sequences = 1.3e-3 average motifs/sequence) and entirely absent from the DHS-based libraries (R0-DHS and R1-DHS), but dramatically enriched in both R1-MPRA (712/1084 = 0.66 motifs/seq) and R2 (568/688 = 0.83 motifs/seq). Several motifs also progressively enriched in DHS- and/or MPRA-based designs include HNF4A/HNF4G, SPIB/ELK1, GATA2/GATA5, and NFE2/JUNB (**Fig. 2E**). The HNF1A and GATA1::TAL1 motifs exhibited significant enrichment in R2 vs both R1-MPRA and R1-DHS, despite no or minimal enrichment in first round design libraries compared to their respective training libraries. These represent motifs with importance emphasized in synthetic rather than genomic enhancers. Overall these results highlight that the models can identify and amplify strong motifs that are scarce in the training data.

Additionally, the synthetic enhancer libraries de-emphasized motifs deemed less relevant to cell type-specific activity. The CTCF motif was significantly depleted in R1-MPRA vs R0-MPRA (3.0e- 3 vs 0.21 motifs/seq); this motif is associated with chromatin organization and is not expected to have differential activity across these cell lines^32^. The SP/KLF motif was significantly depleted in R2 vs R1-MPRA and R1-DHS (3.3e-2 vs 0.76, 0.12 motifs/seq). This motif consists largely of C and/or G repeats, and belongs to a family of universal stripe factors (USFs), which are implicated in promiscuously cooperative activity across many cell types and with many TFBSs^33^. High SP/KLF prevalence in R0-MPRA sequences likely led to strong inclusion in R1-MPRA sequences, but R1 measurements discovered no strong association between these motifs and cell type-specific activity, resulting in R2 depletion.

### Motif analysis elucidates sequence features of highly-specific enhancers

To identify sequence features associated with strong specificity, we first compared motif occurrence in the most specific HepG2- and K562-targeted R2 designs (|log2FC_H2K_| > 3) (**Fig. 2F**). HepG2 enhancers were dominated by TP53 motifs (660 motifs / 395 sequences = 1.67 motifs/seq average), followed by HNF4A/HNF4G (1.06), HNF1A (0.52), HNF4A/NR2F1 (0.43), and TEF (0.29). K562 enhancers were dominated by NFE2/JUNB (247/168 = 1.47 motifs/seq), SPIB/ELK1 (1.38), GATA1::TAL1 (1.07), and GATA2/GATA5 (0.80) (**Fig. 2F**).These findings largely agree with previous reports of cell line-specific TFs^14,23,32,34^.

To dissect the effect of motif multiplicity, we focus on 6 motifs that occur with multiplicity > 1 in more than 50 R2 sequences. For 4 of these we observed a significant dose-response effect between multiplicity and specificity up to 2 motifs (TP53, HNF4A/HNF4G, SPIB/ELK1) or 3+ motifs (GATA1::TAL1), with the marginal effect of adding additional repeats progressively decreasing (**Fig. 2G**). A significant *decrease* in specificity was observed between multiplicities 2 and 3+ for NFE2/JUNB and between 1 and 2 for GATA2/GATA5 (**Fig. S5A**). This effect is possibly explained by additional copies of these motifs decreasing the amount of sequence space available for other, stronger or non-redundant motifs which were not explicitly considered in this analysis; or by off-target activity increasing while on-target activity saturates.

As additional controls in R1-MPRA, we manually designed 62 homotypic enhancers consisting of 1-7 repeats of 9 known TFBS motifs from the CISBP2.0 database embedded in a background of fully random bases (**Methods**). Motifs were selected based on prior reporting of enhancing or repressing roles in HepG2 and K562^14,23^, as well as enrichment analysis on the designed sequences. For TP53, GATA1::TAL1, and NFE2/JUNB, the multiplicity effect was directly confirmed by these manually designed homotypic enhancers, which remove the confounding effect of other motif types (**Fig. S5B**). Saturation occurred at multiplicities 2, 3, and 3 for these motifs, respectively. For NFE2/JUNB, specificity decreased slightly at multiplicities > 3 because even though the on-target cell type activity continued to increase, the off-target cell type activity increased at a greater rate. This motif was deployed at high multiplicity in Max1 designs for both cell types, corroborating this pattern of differential but nonzero activity in both cell lines (**Fig. S5C,D**). One additional motif, HNF1A, exhibited a multiplicity effect (saturation at n=3) in the manual homotypic designs, which was not observed in the global R2 analysis due to low prevalence of high multiplicity sequences. Our findings on motif multiplicity are in good agreement with prior reporting^32,35^. For the remaining 5 motifs for which we tested manual homotypic enhancers, no multiplicity effect was observed (**Fig. S5B**).

Although repeats of the same motif were often associated with increased enhancer specificity, the best enhancers from each library and in each cell type contained more than one motif type. Analysis of the top 5% of HepG2 enhancers in R2 reveals frequent co-occurence of motifs identified by the differential enrichment analysis, with the best enhancers primarily consisting of 1-2 TP53 motifs in combination with HNF4A/HNF4G and/or HNF1A (**Fig. 2H**). The top 5% of K562 enhancers in R2 all contain 2-4 of the following motif types in varying combinations: NFE2/JUNB, SPIB/ELK1, GATA1::TAL1, and GATA2/GATA5 (**Fig. 2H**). We also observe a weak but significant linear association between motif diversity (number of unique motifs) and specificity in both HepG2 and K562-targeted sequences (**Fig. S5E**).

### Motif ablations in top R1 enhancers highlight different modes of motif interaction

In parallel with testing R2 sequences, we also performed perturbation experiments on the original R1 designs to better understand the underlying sequence grammar. Given the high motif density in synthetic enhancers and the possibility of redundant/non-causal motifs, we performed a feature ablation study on the top five R1 enhancers in each cell line. We ablated all single motifs and all pairs by replacing them with randomized nucleotides (**Fig. 3A, Methods**), then experimentally measured enhancer activity. For each single and double ablation, we calculated an ablation score (f_a_) as the fraction of the original enhancer’s activity reduced by the ablation (**Fig. 3B, Methods**) The median single and double ablation scores were 0.13 and 0.30, respectively, validating that the identified motifs generally contributed to enhancer performance, with some ablations achieving much higher effects (max single f_a_ = 0.92, max double f_a_ = 0.93). Because the TP53 motif occurs so frequently in our designs, we were able to ask whether the motif position within the enhancer mattered. Indeed, variation in individual TP53 motif single ablations revealed that TP53 motifs closer to the promoter tended to have higher single ablation scores (**Fig. 3C**).

**Fig. 3.**
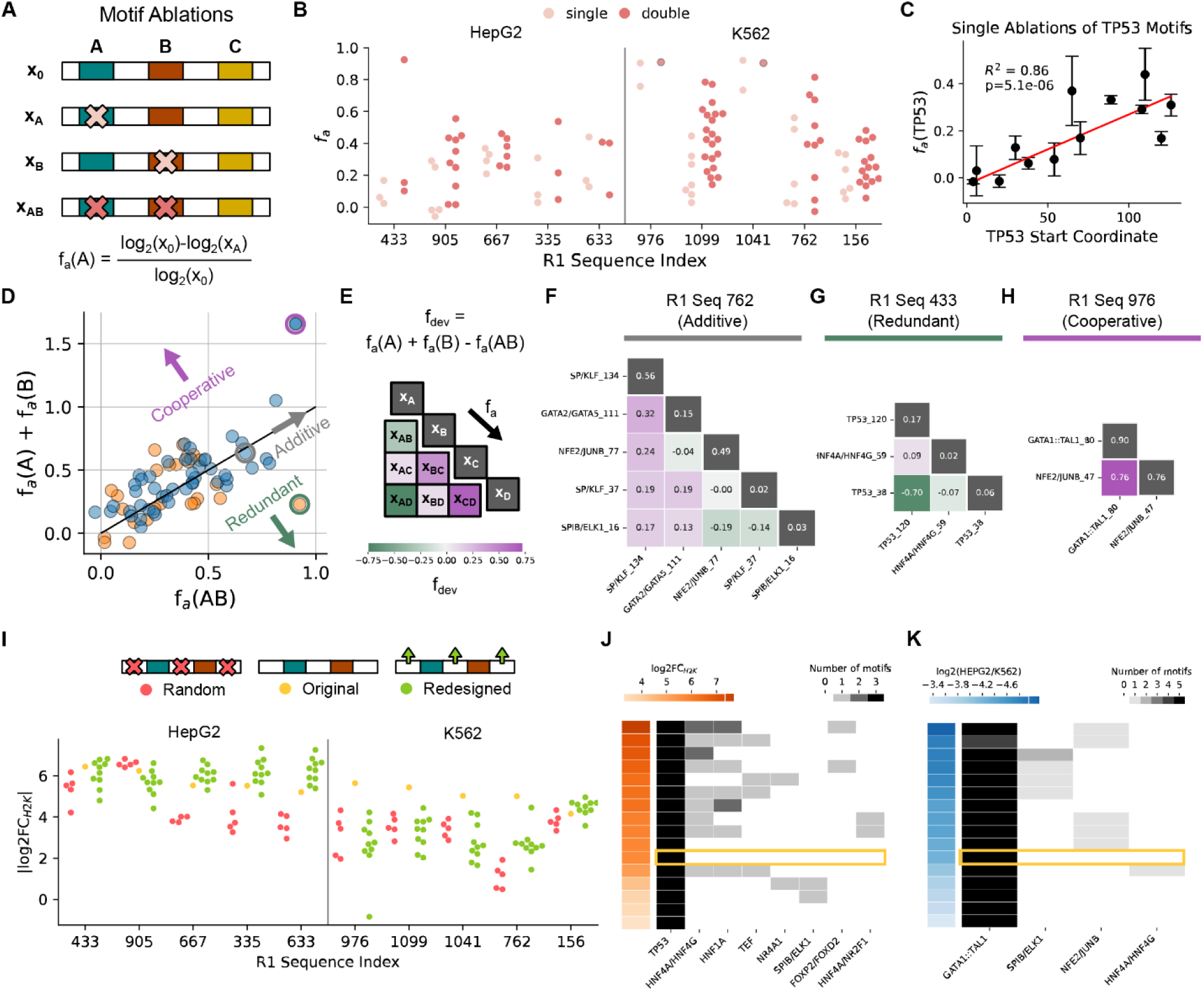
Perturbations of synthetic enhancers indicate causal features of cell type-specific activity. **A,** Overview of single and double motif ablation procedure. Top 5 R1 enhancers in each cell line were perturbed by ablating all individual motifs and motif pairs via replacement with randomized nucleotides. Three randomizations were performed per ablation. Ablation scores were calculated for each single and double motif ablation as the reduction in specificity relative to the unmodified sequence. **B,** Ablation scores for single and double ablations for each R1 sequence. Each point is the mean f_a_ from the three randomizations for a given ablation. For Sequences 976 and 1041 double ablations were not directly measured as these would remove all motifs from the enhancer, and it was assumed this would remove the majority of enhancer activity. **C,** f_a_ regressed on motif position (distance between motif start and sequence start, 0-144) for the eleven single ablations of a TP53 motif. Markers and error bars show the mean and standard deviation from the three randomizations of each ablation. Inverse variance-weighted least- squares regression line shown in red. **D**, An additive model of motif interaction (f_a_(A) + f_a_(B), y-axis) is plotted against the measured double ablation score (f_a_(AB), x-axis). Each point represents an ablated motif pair, and is colored by target cell type of the original enhancer. For R1 Sequences 976 and 1041 the f_a_(AB) was not measured directly, so we estimate with model predictions from M0+1 models fine-tuned on R2 data (**Methods**). Discrepancy between additive model and measurement indicate different modes of motif interaction, as annotated in plot. **E**, Schematic of deviation score calculation and heatmap. Single ablation scores were plotted along the heatmap diagonal, deviation scores for each motif pair were plotted on the off-diagonals. Each heatmap shows deviation scores for every motif pair in a single sequence. Motifs are named by motif type and starting position in the sequence. **F,** Deviation map for R1 Seq 762; most motifs interact additively, e.g. GATA1/GATA5_111 and NFE2/JUNB_77. **G,** Deviation map for R1 Seq 433; contains prominent example of redundancy (TP53_38 and TP53_120). **H,** Deviation map for R1 Seq 976; contains prominent example of cooperativity (GATA1::TAL1_80 and NFE2/JUNB_47). For the double ablation condition, the deviation score was obtained from model predictions. **I**, Top 5 R1 enhancers in each cell line were perturbed by preserving all FIMO-annotated motifs and either 1) randomly shuffling the non-motif sequence, or 2) re-optimizing the non-motif sequence with Fast SeqProp. Swarmplot shows measured |log2FC_H2K_| for these sequence perturbations. **J,K** Motif composition of non-motif redesigns for R1 sequence 335 (**J**), 156 (**K**). Yellow box indicates original sequence, all other rows are redesigns (both Fast SeqProp and shuffled), sorted by descending specificity.

Finally, we tested an additive model of motif interactions, comparing each double ablation score to the sum of the corresponding single ablation scores, and defining the difference as the deviation score, f_dev_ (**Fig. 3D-E, Fig. S6, Methods**). The majority of motif interactions were additive (f_dev_ ≅ 0), indicating motifs had independent activity/contribution to enhancer strength (e.g. R1 Seq 1099, **Fig. 3F**). R1 Seq 433, which contained two TP53 and one HNF4A motifs, exhibited redundancy (f_dev_ << 0), wherein ablation of either of its two TP53 motifs resulted in only mild reduction of enhancer activity, but ablation of both TP53s knocked it out almost completely (**Fig. 3G**). This sequence can be compared to R1 Seq 633, which consists of three TP53 motifs, where no single or double ablation achieved more than moderate impact on enhancer activity (**Fig. S6F**). In contrast, from the R1 Seq 976 and 1041 ablations we uncovered a prominent example of cooperativity (f_dev_ >> 0) between the GATA1::TAL1 and NFE2/JUNB motifs. Ablating either of these motifs reduces most of the enhancer activity, indicating both are necessary for the function of this enhancer (**Fig. 3H, Fig. S6G**).

### Optimizing non-motif sequence in top R1 enhancers can improve specificity

We next investigated whether we could improve enhancer specificity by redesigning sequences outside the identified motifs. We selected the top five R1 enhancers in each cell line and re-optimized all bases outside of an aligned motif with Fast SeqProp and the M0+1 model, generating 10 new designs per original enhancer. As controls, we generated 3 sequences where motif flanks were replaced with random bases. Re-optimizing non-motif sequence improved specificity for 5/5 HepG2 designs and 1/5 K562 designs (**Fig. 3I, Fig. S7**). The majority of HepG2 designs were improved by embedding more non-TP53 motifs (e.g. **Fig. 3J**), indicating initial designs reached saturation of TP53 multiplicity effects (**Fig. 2G, Fig. S5E**). The most successful HepG2 redesign (log2FC_H2K_ = 7.35) was one of the most specific enhancers even compared to R2, representing a 1.33X improvement over the original sequence. The successful K562 re-design originally consisted only of GATA1::TAL1 repeats, and was improved by adding either a SPIB/ELK1 or NFE2/JUNB motif, the latter completing the combinatorial pair implicated by ablation analysis (**Fig. 3K**); the best redesign achieved ∼1.73X improvement over the original enhancer. In general, K562 re-designs were likely less successful because of lower model accuracy on the K562 prediction task as well as higher motif density in the original sequences, resulting in less “free” sequence to re-design. Notably, randomizing non-motif sequence significantly, if modestly, reduced specificity for 9/10 tested enhancers compared to the original sequence (**Fig. 3I**).

### Shorter enhancers display high specificity

To further refine our understanding of the core functional grammar of enhancers, we attempted to reduce enhancer size by designing 72, 50, and 25bp-long sequences. In both cell lines, shorter enhancers down to 50bp achieved statistically equivalent activity to those with a length of 145bp by Wilcoxon rank-sum (**Fig. 4A**). In fact, the best 72bp HepG2 enhancer had statistically equivalent specificity to that of the best 145bp HepG2 enhancer **(Fig. 4B)**. A significant decrease was only observed at 25bp (HepG2: 3.62 vs 5.80, K562: 0.15 vs −3.08). In HepG2 designs, 25bp enhancers consisted of a single TP53 motif, which was sufficient to drive moderately specific activity. Conversely, 25bp K562 designs consisted of a single GATA1::TAL1 motif which could not drive enhancer activity alone, in agreement with the ablation analysis. As sequence length is increased, HNF4A/HNF4G and HNF1A motifs are the next motifs to be prioritized after TP53 in HepG2 enhancers, whereas NFE2/JUNB and SPIB/ELK1 are prioritized after GATA1::TAL1 for K562 (**Fig. 4C**).

**Fig. 4.**
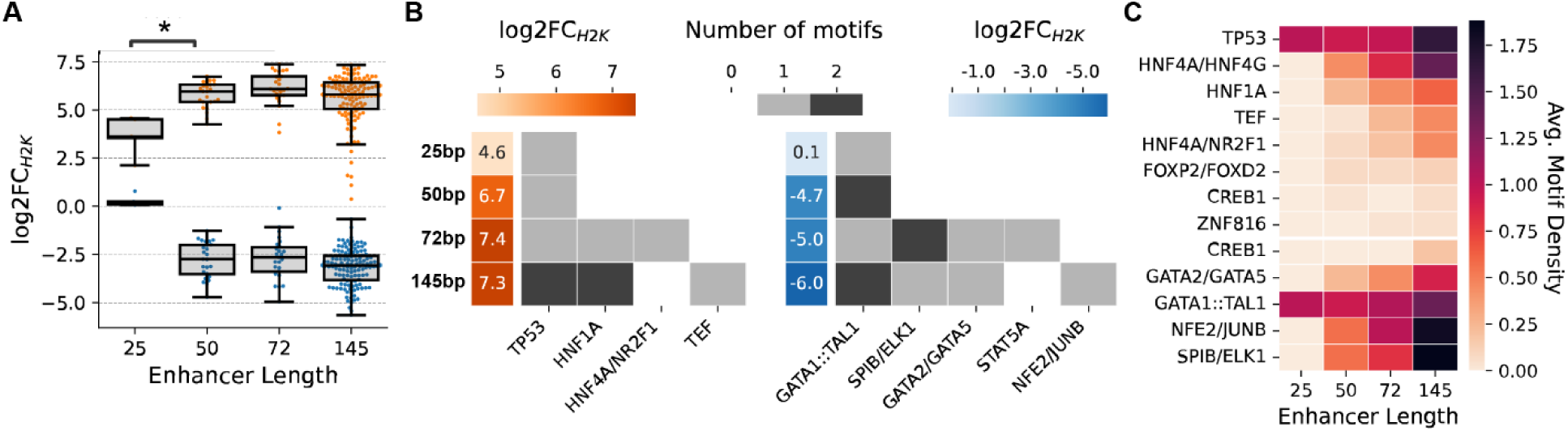
Sorter enhancers display high specificity. **A,** Boxswarm plots show log2FC_H2K_ for R2 enhancer designs grouped by sequence length. Asterisks indicate significant difference in median specificity by Wilcoxon rank-sum (p < 0.05). **B**, Motif content and log2FC_H2K_ of most specific enhancer of each sequence length for HepG2 designs (left) and K562 designs (right). Each row corresponds to a different enhancer length, denoted by text annotation to the left of plot. log2FC_H2K_ colormaps computed separately within each cell type. **C,** Average motif density of R2 enhancers as a function of sequence length shown for motifs with average motif density >=0.05 at every sequence length. Horizontal white line separates HepG2- from K562- targeted enhancers, average motif density calculated separately within each cell type. Motifs sorted by enrichment in 145bp enhancers (descending in HepG2, ascending in K562).

### A scMPRA enables characterization of enhancer activity at the single cell level

Finally, we sought to characterize cell-to-cell heterogeneity in enhancer activity and understand how it may depend on cell state and TF expression. To this end, we performed an scMPRA following an approach similar to Zhao *et al.*^10^ (**Fig. 5A**, **Methods**). Briefly, we modified our R1-MPRA library by incorporating a random barcode and transfected it into a 1:1 mixture of K562 and HepG2 cells, followed by combinatorial split-pool indexing to tag mRNAs with cell barcodes^36^ and dual sequencing of transcriptome and MPRA library cDNAs (**Fig. S8A**). We recovered 1,343 enhancers across 10,640 cells at a median of 4 unique enhancers per cell (**Fig. S8B, C**). We verified that our previous bulk MPRA measurements could be accurately reconstructed from pseudobulk analysis of scMPRA data (Pearson R = 0.91, **Fig. 5B**). Subsampling cells shows that such high correlation is robust down to pseudobulk size of ∼1,000 cells (**Fig. S8D**). Thresholding by minimum expression levels in each subsample maintains the high correlation, indicating that the sampling noise is due to dropout events attributable for example to transfection efficiencies or mRNA capture sensitivities. To understand whether our synthetic enhancers show unexpected cell state specificity, we examined cell cycle phases and a known major “stem-like” differentiation/proliferation state within K562 cells marked by CD24 expression^37^. Examining differential pseudobulk enhancer activities, we found that our synthetic enhancers are largely robust across these substates (**Fig. S8H-O**).

**Fig. 5.**
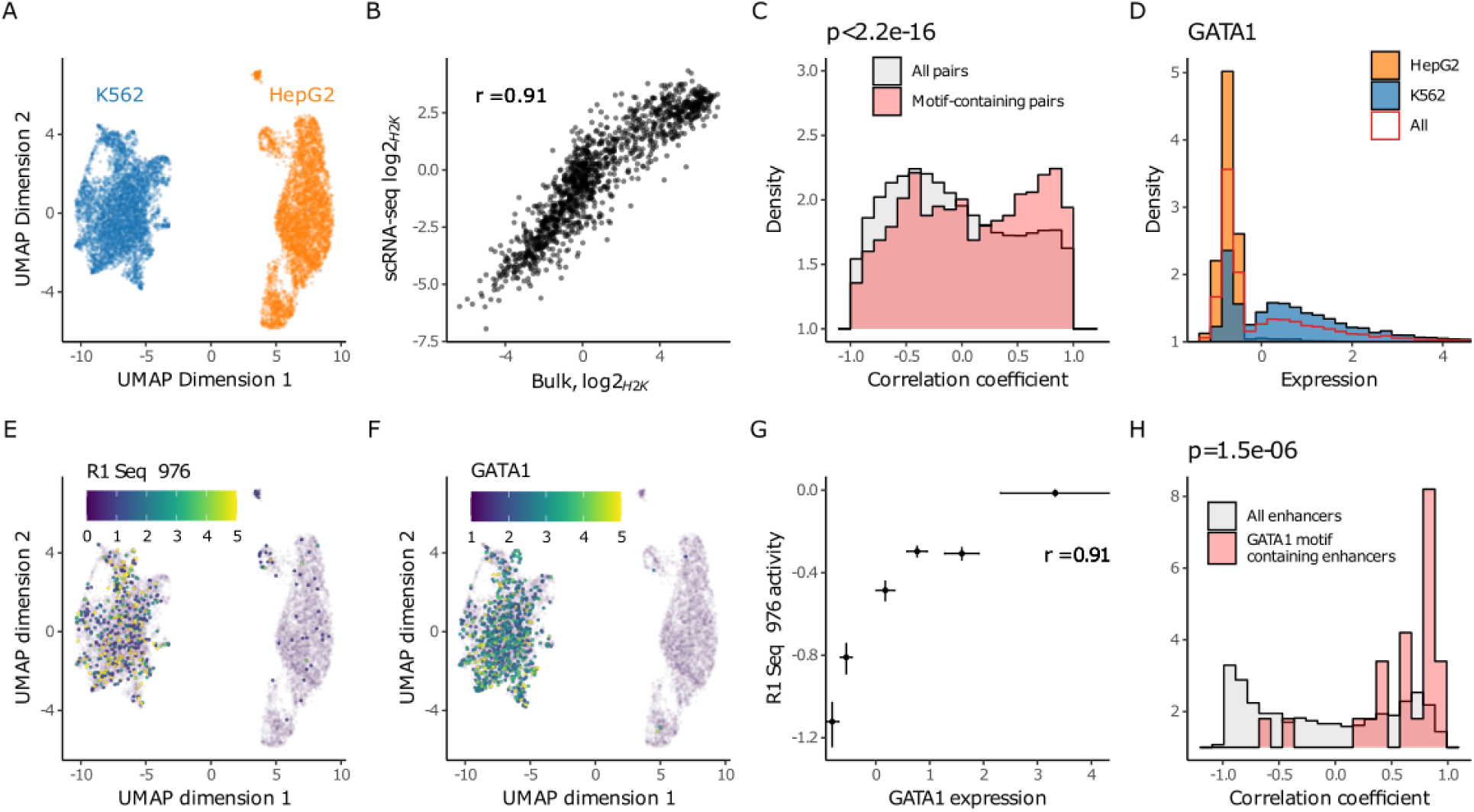
Synthetic enhancer activity confirmed at the single cell level. **A,** UMAP projection of the single cell transcriptomes. Clusters corresponding to K562 and HepG2 cell types are colored in blue and orange respectively. **B,** Comparison of cell-type specific activity of synthetic enhancers obtained by bulk (x axis, same as Fig. 1B) vs. single-cell analysis (y axis), where the activity was calculated via pseudobulk aggregation of the clusters shown in panel A. **C,** Distribution of all possible pairwise correlations between transcription factors and enhancer expression across pseudobulk of single cells binned by transcription factor expression values. See panel G for an example of one such pair. In red are pairings where the enhancer contains the DNA sequence motif for the transcription factor. p value from two-sided Kolmogorov-Smirnov test. **D,** Expression levels of GATA1 transcription factor across: all cells in red; HepG2 in orange, K562 in blue. **E,** Activities of R1 Seq 976, containing GATA1 transcription factor motif, across single cells atop the UMAP projection of the transcriptomes. **F,** Expression levels of GATA1 transcription factors across single cells atop the UMAP projection of the transcriptomes. **G,** Pseudobulk correlation of GATA1 transcription factor expression levels and R1 Seq 976. The pseudobulks are binned by GATA1 expression levels. Vertical error bars represent bootstrap standard errors of enhancer activities. Horizontal error bars represent standard deviation of the transcription factor expression levels across the cells in each pseudobulk bin. **H,** Distribution of correlations between GATA1 expression versus all enhancers across pseudobulk of single cells binned by GATA1 expression values. In red are pairings where the enhancer contains the DNA sequence motif for GATA1. p value from two-sided Kolmogorov-Smirnov test. Also see Supplementary Fig. 10.

If the TFBS motifs in our synthetic enhancers are indeed causal, we would expect to observe a relationship between individual TF expression and enhancer activities. To investigate this hypothesis, we binned cells by the expression levels of a given TF and obtained per-bin pseudobulk MPRA reporter expression levels for each enhancer. We then calculated a correlation value for every TF-enhancer pairing across all bins. We generally observe positive correlations between pairs of enhancers and TFs if the enhancer contains a motif corresponding to the TF in the pairing. In contrast, TF-enhancer pairs where the enhancer does not contain the corresponding TFBS show generally lower and negative correlation coefficients (**Fig. 5C**). For example, R1 Seq 976 is K562-specific and contains one GATA1::TAL1 binding site (**Fig. S9A**). The scMPRA recapitulates its K562-specific activity (**Fig. 5D,E**), along with the expected GATA1 expression pattern (**Fig. 5F**). We find a positive correlation between reporter levels from this enhancer and GATA1 expression across the bins (**Fig. 5G**). We observed similar positive correlations for other GATA1:TAL1 motif-containing enhancers, but not when considering other enhancers without GATA1:TAL1 motif against GATA1 expression (**Fig. 5H**). Similarly, R1 Seq 370 is HepG2-specific and contains a TP53 TFBS (**Fig. S9B**). Its activity is correlated with TP53 expression at the single cell level, as are the activities of other TP53-containing enhancers (**Fig. S9C-G**). Thus, cell type specific enhancer activities are predicted by levels of TFs that are differentially expressed across cells.

## Discussion

In this work we demonstrate iterative ML-based design and experimental validation of functional, cell type-specific enhancers in human cell lines. Across two generations of sequence design we achieve enhancers that drive cell type-specific gene expression more strongly and with a higher relative success rate than putative enhancers sourced from accessible regions of the genome, either selected directly from accessibility measurements (R0-DHS) or screened via MPRA (R0-MPRA). We also find that sequences generated in a second round of design were more specific than those from the first round despite only a relatively small number of sequences being available for model retraining. Our models are able to both interpolate (design for intermediate levels of differential expression) and extrapolate (design beyond the range of measured expression strengths), via *ab initio* discovery and implementation of a condensed TFBS motif grammar.

Analysis of synthetic enhancers and corresponding followup experiments implicated several sequence features driving stronger specificity compared to genome-sourced enhancers: most prominently, higher motif density coupled with drastic enrichment of a concise vocabulary of TFBS motifs with cell type-specific activity. While top-performing enhancer designs featured motif repeats, they were also heterotypic, suggesting that deep learning models capture an impact of motif diversity for optimizing sequence specificity.

The success of model-designed enhancers implementing a relatively small set of TFBS motifs may erroneously suggest the possibility of a simpler, model-free design strategy: manual embedding of relevant motifs identified by enrichment analysis in putative enhancers and/or accessible regions of the genome. However, this identification is nontrivial. The TP53 motif was not enriched in our training data, but was discovered to be the strongest driver of HepG2- specificity in our synthetic libraries. TP53 has previously been implicated as a pioneer factor capable of binding to and initiating the opening of closed chromatin, which may explain its infrequency in training data derived from accessible regions of the genome^39^. We note that another recent MPRA in both HepG2 and K562 cells, which reports testing a “nearly comprehensive set of all annotated CREs” in these cell lines (including 66,017 putative HepG2 enhancers), associates HNF4A but not TP53 with HepG2 enhancer activity^34^, in line with our findings based on DHS data (R1-DHS). Even when prior reporting of TF function is available, the complex context dependencies of the cis-regulatory code mean TFs may still behave unexpectedly in different cell types. As an example, the NFE2/JUNB motif was discovered to be enriched in the top K562-specific enhancers of our designs, and exhibited a positive association between multiplicity and differential enhancement strength. However, this motif corresponds to a TF described as having non-specific “housekeeping” activity^34^, and was not identified in two distinct papers studying the Sharpr-MPRA dataset^14,23^, or the recent MPRA by Agarwat *et al.* as associated with K562-specific enhancement^34^; nor was it prevalent in the DHS-based designs.

A scMPRA supports a relationship between TF mRNA abundance and enhancer activity. However, while these data are consistent with a causal relationship between TF expression and the activity of enhancers with cognate TFBSs, there are several caveats. First, TF mRNA levels may not be predictive of TF activity. For example, while enhancers containing the TP53 motif have no activity in K562 cell lines, TP53 mRNA levels are only two-fold lower in K562 than HepG2. However, previous work found that both TP53 alleles in K562 encode a non-functional protein, but the relevant sequence variation is not captured in a typical scRNA-seq workflow^38^. More generally, the activity and localization of TFs is often controlled through post-transcriptional modification (e.g. phosphorylation) not captured by RNA-seq. Moreover, we find that for many TFs, expression differences between cell types are larger than expression variation within each cell type. As a result, observed correlations may be driven primarily by the expression differences between the two cell types, resulting in spurious correlations due to the fixed activity ratio between different TFs at the bulk level. Nevertheless, we expect that given its capability to measure full pairwise variations in enhancer activities and TF expression levels, future scMPRAs on heterogeneous cell populations or in combination with multiplexed knockdown/overexpression perturbations promises an effective experimental approach to infer causal relationships between enhancers and TFs in a highly parallelized, pooled manner.

Highly specific enhancers were generated with disparate design strategies, and we find evidence that model accuracy is more performance-limiting than choice of design algorithm. Notably, we achieved successful designs using models trained on a noisy dataset of enhancement strength measurements (R0, ∼30K sequences, average read count inter-replicate Spearman R = 0.61); as well as using models trained only on the smaller R1 dataset (∼1.3K sequences, average read count inter-replicate Spearman R = 0.99). In fact, the best K562 enhancer in R2 was designed using M1 models (log2FC_H2K_ = −5.99)

As a training set the R1 libraries differ from R0 along several axes: the sequences are primarily model-generated, exhibit higher motif density, have a greater proportion of strong and specific enhancers (i.e. more “positive” class balance), and have less noisy measurements. While data quality may be an important factor, we hypothesize that the abundance of strong, functional examples in R1 libraries make them more efficient datasets for model training. A recent study from the Cohen lab used active learning, where one explicitly selects samples to maximize model uncertainty, to refine a model of CREs over 4 design-experiment cycles^40^. While their approach required estimating model prediction uncertainty, our results suggest that simply designing sequences with extreme behavior may result in high quality training datasets. Future work might more rigorously explore which dataset features are most important for model training; nonetheless, these results indicate the potential for small, low-cost synthetic libraries to enable expansion of enhancer design into new cell types.

For the first round of designs (R1) we pursued two complementary approaches. A first approach, R1-MPRA took advantage of published MPRA data in the target cell lines, similar to a recent study^22^. Given that MPRA data is only available for a limited number of cell types and tissues, we additionally explored training models directly on chromatin accessibility, effectively bypassing the requirement for MPRA data. Our results show that this is a feasible approach, yielding functional enhancers in both HepG2 and K562. However, accessibility-based enhancer activities were on average lower and fewer sequences were strongly specific compared to those designed from MPRA-trained models (**Fig. 1F**). The use of a GAN for the DHS-based designs, which explicitly regularizes sequences to be similar to the (accessible, but not necessarily highly specific or active) genomic sequences used in training, may have also impacted performance. Nevertheless, the broad availability of chromatin accessibility data opens up opportunities for future work evaluating the generalizability of this approach to more diverse cell types, as well as strategies to bring performance on par with MPRA-based models with no or very little reporter measurement data.

An important caveat to comparisons with genomic sequences in this and other work^22^ is the confounder of sequence length. Natural enhancers are believed to reach up to kilobase lengths^2^, and excerpting only small windows of genomic sequence likely captures many incomplete enhancers, which could theoretically disadvantage them against synthetic enhancers explicitly designed to be <145 bp. Accordingly, we find a relatively low motif density in both R0 libraries, especially when compared to our designed enhancers (**Fig. 2A**). Thus, a “fair” comparison to genomic enhancers might require large DNA fragment synthesis and lower throughput experiments. However, ∼1kb genome-sourced enhancers have been characterized in some contexts, and even then found to have a relatively low success rate^5,11,41^. Furthermore, designing shorter functional enhancers is useful for gene therapy applications, where AAV vectors are limited by the size of their payload^42^. Here, we show that enhancers can be shortened to 50 bp without significant reduction in strength, suggesting core functional grammar can be greatly compressed in directly promoter-adjacent enhancers.

Finally, while the success of our designed library is encouraging, we acknowledge that achieving cell type-specificity in two cell lines is a starting point, not the end goal. The more interesting, biologically relevant challenge will entail designing specific enhancers for a larger panel of cell types, including closely related ones. In this situation, individually strong TFs without crossover activity in multiple cell types may be more difficult to identify, and combinatorial syntax taking advantage of individually weaker but more specific TFs could become more important. A deep learning approach should be the most readily extensible to this task, and theoretically enable precise tuning of activity in arbitrary sets of cell types via intelligent, efficient exploration of the full combinatorial space of motifs, relative motif positioning, and non-motif flanking sequence–i.e., the complete cis-regulatory code. While the dearth of functional MPRAs beyond a narrow range of well-studied cell lines presents an obstacle to more widespread expansion of this approach, our success with accessibility-based designs suggests a path forward. Together, our approach and current results provide a foundation for ongoing and future studies to the data-driven design and understanding of synthetic regulatory elements.

## Acknowledgements

This work was supported by NSF award EF-2021552 and NIH award 1R01HG012346 to GS and NIH NHGRI award R35HG011317 to WM.

## Author contributions

Conceptualization, C.Y., S.C., G.S., W.M.; Methodology, C.Y., S.C., G.W.B., P.B., G.S., W.M.; Software, C.Y., G.W.B., P.B.; Formal Analysis, C.Y., S.C., G.W.B., P.B., W.M.; Investigation, C.Y., S.C., G.W.B., P.B., W.M.; Resources, G.S. and W.M.; Writing - Original Draft, C.Y., S.C., G.W.B., G.S., W.M.; Supervision, G.S. and W.M.

## Supplementary Information

Document S1. Figures S1-11

Document S2. Tables S1-5

## Methods

### Preparation of filtered Sharpr-MPRA (R0) for R1-MPRA model training

We downloaded the raw read count-level Sharpr-MPRA dataset from Gene Expression Omnibus, accession number GSE71279, via the Dropbox hosting link provided by Movva et al.: https://www.dropbox.com/sh/wh7b30dauxuajcw/AABQsvfmG65knGbFv0UsIcv1a?dl=0. The Sharpr-MPRA library contained a few duplicated sequences, thus measurements for 4912 enhancers are included in the dataset 2-3 times. These were not explicitly removed for the initial model training. We filtered out all sequences with a minP DNA read depth < 200, log2(HEPG2) <-2.4, or log2(K562) < −2.4 in either replicate. log2(RNA/DNA) were calculated using raw RNA and DNA counts with a pseudocount of 1.

### R1-MPRA model training

The model architecture used in generating R1-MPRA is constructed as follows. The one hot-encoded Input Layer (145x4) is passed in parallel to three branches of convolutional layers. Each branch contains 600 filters of size 7, 11, or 25; and feeds the output of the convolution through BatchNormalization, ReLU activation, Global MaxPooling, and Dropout (0.075). The outputs of each branch are concatenated and passed to a Dense layer with 64 nodes and ReLU activation, before finally feeding into the 2-node output layer with linear activation predicting log2(HEPG2) and log2(K562). Models were trained in TensorFlow using the ADAM optimizer (learning rate = 2e-4, beta_1 = 0.9, beta_2 = 0.999) and a batch size of 64, for 60 epochs with early stopping (patience = 8, min_delta = 1e-6). All data splits were augmented with reverse complements.

To avoid overreliance on any single model for sequence design, we trained a total of 120 models divided in three categories. First, we trained 10 “Single” models on the same random training/validation/test data split but with different randomly-initialized network weights. Second, we trained 100 CNN models with the same architecture on randomly resampled bootstraps of the merged training and validation data splits (“Boot” models). Third, we formed 10 ensemble predictors by splitting the bootstrap models into groups of 10 and averaging the outputs of every model within the group (“Ensemble” models). Evaluated on the same held-out test dataset, prediction-measurement correlations from all models were on par with inter-replicate correlations (**Fig. S1C**), with Ensemble models having the best performance, followed by Single and Boot models.

### R1 design method implementation

For the R1-MPRA library, we used three design methods based on different general principles: simulated annealing (Monte Carlo sampling), Fast SeqProp (gradient descent) and Deep Exploration Networks (DENs) (deep generative models). Sequences were designed to maximize or minimize log2FC_H2K_, clipped to a threshold.

### Deep Exploration Networks

In Deep Exploration Networks^25^ a generative neural network is trained to produce sequences that simultaneously maximize a fitness score derived from a predictor model while penalizing the similarity of generated sequences. At a high level, the cost function for training a DEN has 3 components: a fitness cost, diversity cost, and entropy cost, each with an associated weight. Here, the fitness cost was max(log2FC_K562_ - log2FC_HepG2_ + fitness_target,0) for HEPG2-targeted designs, and max(log2FC_HepG2_ - log2FC_K562_ + fitness_target,0) for K562-targeted designs. This loss function was implemented because empirically it was found that maximizing the predicted model output unbounded yielded sequences with unnatural characteristics that we initially hypothesized to be undesirable. Without this loss clipping, generated sequences were predicted to have cell type-specificity several orders of magnitude beyond the training data, and exhibited low diversity due to strong single or di-nucleotide repeats. The following parameters were used in the cost function for training all DENs: fitness target = 2.5, fitness weight = 0.0075, entropy_min_bits = 1.8, entropy weight = 0.5, similarity margin = 0.5, similarity weight = 5.0.

For each generator trained, we generated 1000 sequences and took the top 10 highest HEPG2- or K562-specific predicted sequences. DENs were used to generate 10 sequences per model per cell type from Single models 0-9, Boot models 0-9, and Ensemble models 0-9 (600 sequences total).

### Fast SeqProp

Fast SeqProp^24^ is a technique previously developed by our lab in which a PWM is optimized via gradient descent in conjunction with the straight-through gradient estimator, such that sequences sampled from the PWM maximize the output of a predictor model. For each sequence generated with Fast SeqProp, we ran the algorithm for 200 gradient updates to generate 10 sequences, then included only the sequence with the highest (HepG2 target) or lowest (K562 target) log2FC_H2K_ score in our designed library. We use n_sequences=10, n_samples=1, n_epochs=1, steps_per_epoch=200, pwm_target_bits=1.8, pwm_entropy_weight=0, fitness_target=3, and ADAM optimizer with learning_rate=1e-3, beta_1=0.9, and beta_2=0.999. The loss function used is max(log2FC_K562_ - log2FC_HepG2_ + fitness_target,0) for HEPG2-targeted designs, and max(log2FC_HepG2_ - log2FC_K562_ + fitness_target,0) for K562-targeted designs. Fast SeqProp was used to generate 2 HEPG2-targeted sequences and 1 K562-targeted sequence from Single models 0-9 (2 K562 sequences per model intended, the 2nd omitted by error), and 1 sequence per model per cell type for Boot models 1-99 (model 0 omitted by error). 228 sequences total.

### Simulated Annealing

In simulated annealing^44,45^, sequences are randomly initialized, then at each step a random mutation is introduced at a random position and probabilistically accepted based on the current temperature value and the change in predicted fitness score. The temperature value is decayed with each iteration so that the algorithm is less likely to accept deleterious mutations as the step number increases. Here, we run simulated annealing for 1000 iterations per sequence, decaying the temperature from an initial value of 0.1 to a minimum value of 0.05 after every 100 iterations, using an exponential scale factor of 0.143. The loss function minimized was log2FC_K562_ - log2FC_HepG2_ for HepG2 targets, and log2FC_HepG2_ - log2FC_K562_ for K562 targets. Simulated Annealing was used to generate 1 sequence per model per cell type for Single models 0-9 and Boot models 1-99 (model 0 omitted by error). 218 sequences total.

### Hand-crafted motif repeats

Handcrafted homotypic motif repeat enhancers were designed for 9 known TFBS motifs from the CISBP2.0 database, which were embedded in randomly generated sequences with multiplicity values of 1-7 for all motifs except HNF4A (multiplicity=1 omitted by error). 62 sequences total. Motifs were selected based on prior reporting of HepG2 and K562 specificity, as well as enrichment analysis on the designed sequences (**Table S3**).

Random sequences were generated using the following background distribution from R0: A=0.230, C=0.254, G=0.284. T=0.231. Each sequence used a different random background.

### R0 control sequence selection

In total 100 sequences from the filtered Sharpr-MPRA dataset were re-measured in R1: 25 selected from both the 99th and 1st percentiles of log2FC_H2K_ scores, and 50 selected at uniform intervals along the log2FC_H2K_ range. The reverse complements of these 100 control sequences were synthesized as well.

### R1-DHS design and model training

#### Generative Adversarial Network (GAN) training

We use a training set of 758k DHSs located on chromosomes 3-X to train a Generative Adversarial Network (GAN)^26^ for generating synthetic sequences with characteristics similar to endogenous human accessible genomic regions. Specifically, we obtained delineations and annotations for 3.5M+ DNase I Hypersensitive Sites (DHSs)^8^, and filtered out DHSs with length less than 200bp and an annotated “mean signal” confidence score less than 0.5. We truncated each DHS to 200bp, centered on their annotated “centroid” position. In case a DHS summit position is less than 100bp from its annotated start or end position, we compensate by including additional length at the other end of the summit. This results in a set of 918,057 DHS sequences of length 200bp. We split these into general training (758,692 sequences, 82% of total, chromosomes 3-X), validation (78,108, 9%, chromosome 2) and test (81,257, 9%, chromosome 1) sets.

The GAN model consists of basic generator and discriminator functions using both convolutional and fully connected layers (**Table S4**). To improve training stability and maintain diversity of generated sequences, we use hinge loss and apply spectral normalization^46^ to the convolutional and fully connected layers of the discriminator. After training, the generator provides a mapping from random input seeds (length 100) to synthetic DNA sequences (length 200bp). Generated sequences generally reflect characteristics of endogenously accessible sequences, as shown by matching nucleotide composition of G/C content and 4-mer sequence patterns (**Fig. S1A,B**).

### Classifier training

We tune generated sequences *in silico* to increase specificity to cellular contexts of interest by way of a separately trained multi-class classification model (**Figure 1D, Table S5**) discriminating between three classes of accessible elements: 1) ‘K562’, 2) ‘HepG2’ & 3) ‘Other’.

These classes are defined using a combination of DHS component annotations^8^ and observed accessibility in K562 or HepG2 cells. To maximize the amount of component-relevant signal, minimize inclusion of experimental noise and promote a high-contrast classification task, we select DHSs for each class to be relatively component-selective, as follows. We define the “purity” of each DHS as the proportion of its dominant component annotation divided by the sum of all component annotations. We then define the ‘Other’ class of DHSs by selecting the 625 highest purity DHSs from each of 16 NMF components, excluding any DHSs that occur in HepG2 or K562 biosamples, for a total of 10k DHSs. For the other two classes, we focus on the ‘Digestive’ component for HepG2 and the ‘Myeloid/Erythroid’ component for K562. Specifically, we co-rank Digestive-component purity and the number of HepG2 biosamples in which a DHS is accessible to select the top ranking 10k ‘HepG2’ DHSs. We follow the same procedure for the ‘K562’ class, using the ‘Myeloid/erythroid’ component. We apply this procedure to chromosomes 3-X to create a dataset of 30k DHSs with high-confidence component labelings, to be used as a training set. We follow the same procedure to select 1k sequences per class from chromosome 2 as a validation set for tuning model hyperparameters, and similarly from chromosome 1 as a test set.

### Sequence tuning

We use the output nodes of our trained classification model to guide the sequence tuning process. Specifically, we search the latent space of the trained generator network for sequences that maximally activate the classification node corresponding to the class of interest. For each tuning run, one class is chosen as a target, and the final classification node (pre-softmax) corresponding to that class is multiplied by −1 and used as loss. We then backpropagate this loss through the networks to the generator’s latent space, and update the input seed to generate a slightly more context-specific sequence. For each generated sequence, we perform 10,000 tuning iterations, converging to an input seed that results in high activation of our target class.

To generate a tuned set of sequences for subsequent experimental validation (R1-DHS), we generated and tuned 300 sequences to the K562 and HepG2 target classes. For each sequence and each target class, we selected a single iteration after convergence as the final representative tuned sequence for subsequent experimental validation. The median selected iterations were 4750 (HepG2) and 3010 (K562), well within range of any notable sequence divergence relative to the training set (**Fig. S1E,F**), yielding sequences that are primed for accessibility in the selected cellular contexts, while preserving identity with the originally generated untuned sequence. We refer to this library as R1-DHS.

### Motif calling

For all libraries (R0, R0-DHS, R1, R1-DHS, and R2), we separately ran FIMO with the default p-value threshold of 1e-4, scanning sequences for matches to the JASPAR 2022 Non-redundant Vertebrate database. We then applied an additional q-value filter, discarding all motif hits with q-values exceeding a threshold of 5e-2. Finally, we applied a custom position-wise clustering algorithm to collapse overlapping motif hits. First, within each sequence we collapsed all overlapping motifs with the same motif ID to the instance with the lowest p-value. Then, we scanned across all the positions in a sequence from 5’ to 3’ and for any position included in multiple motif hits, we kept the motif hit with the lowest p-value and discarded the rest. We allow an overlap of 3 bp without collapsing motifs together.

For subsequent analysis, we additionally clustered motifs using the JASPAR Core Vertebrates RSAT clusters (841 motifs, 137 clusters). (<u>JASPAR - JASPAR CORE Vertebrates clustering</u> <u>(genereg.net)</u>).

For the purpose of Fig. S2G,H, we obtained motif files `HNF4A_nuclearreceptor_3` and `SPI1_ETS_1` from https://www.vierstra.org/resources/motif_clustering and scanned sequences using FIMO with a threshold of 1e-5.

### Second generation model training

To train models for R2 enhancer design, R0 data was reprocessed with DESeq2, sorted by descending log2FC_H2K_, then split into 10 crossfolds snakestyle to ensure equivalent distributions of enhancement strengths in all the folds. R1-MPRA and R1-DHS were split into crossfolds according to the same scheme, with only 5 crossfolds used for R1-DHS due to the smaller library size.

We pre-trained 9 independent models (“M0”) on the DESeq2-processed R0 splits, holding out Crossfold 0 as a constant test set and rotating through the remaining crossfolds as the validation set. Before retraining, we added L2 regularization to the dense layer weights after model interpretation indicated overemphasis on a single TP53 motif per sequence even in sequences with high TP53 multiplicity; while this may be an accurate representation of the biology, we deemed this more likely a model pathology that might also obscure the importance of non-TP53 motifs. We then re-optimized model hyperparameters using the keras_tuner implementation of the Hyperband strategy, resulting in minor tweaks to layer and filter sizes. The re-optimized architecture consisted of 3 parallel branches of 608 filters each of size 11, 15, and 21, trained with a dropout rate of 0.181; followed by a dense layer of size 224 with an L2 weight of 1e-3; trained with the Adam optimizer (learning rate = 3e-4, beta_1 = 0.9, beta_2 = 0.999) and early stopping monitoring the validation loss (min_delta = 1e-6, patience = 5). To accommodate eventual training on R1-DHS data, the input layer was expanded to size 200; any sequences shorter than this were 0-padded symmetrically on both sides.

To train M0+1 models, we held out Crossfold 0 from the R1 and R1-DHS datasets, then finetuned each M0 model on the remaining data, rotating through the R1 crossfolds as the validation set; all R1-DHS splits were used for training. A learning rate of 1e-4 was used, in addition to a ReduceLROnPlateau callback (factor = 0.25, min_delta = 1e-2, patience = 4). Models were finetuned for a maximum of 250 epochs using early stopping (min_delta = 1e-3, patience = 5), batch_size = 64.

Finally, we trained 9 M1 models using the same architecture and optimizer hyperparameters as M0 models, but using only the R1 and R1+DHS splits without a pre-training step with R0.

### Model interpretation

M0+1 ensemble model predictions were interpreted on all R1-MPRA sequences using the DeepExplainer library of the SHAP^47^ package, 100 dinucleotide shuffles per sequence. SHAP values were obtained for both log2FC_HepG2_ and log2FC_K562_ predictions.

### R2 Design

All R2 designs were Fast SeqProp-based, and implemented an additional loss term not used for R1 designs (pwm_loss) penalizing single nucleotide repeats. We used target_weight = 1, pwm_weight = 2.5, entropy_weight = 1e-3, learning_rate = 1e-3, n_iter_max = 1000, and default for all other parameters.

### Unbounded objective

An unbounded Fast SeqProp objective was used with both M0+1 and M1 models. 200 sequences were designed to maximize both HepG2- and K562-specificity unbounded, and the top 130 (M0+1) or 105 (M1) predicted sequences for each cell type were selected. The loss function minimized was log2FC_K562_ - log2FC_HepG2_ for HEPG2-targeted designs, and log2FC_HepG2_ - log2FC_K562_ for K562-targeted designs. 470 sequences total.

### Clipped objective

200 sequences were designed to maximize both HepG2- and K562-specificity with a clipped Fast SeqProp objective, and the top 110 predicted sequences for each cell type were selected, using the M0+1 model. The loss function minimized was log2FC_K562_ - log2FC_HepG2_ - target for HEPG2- targeted designs, and log2FC_HepG2_ - log2FC_K562_ - target for K562-targeted designs, where target = 1.1X the highest predicted specificity on R1 sequences according to the M0+1 model. 220 sequences total.

### Max1 and Min1 designs

100 sequences each were designed to maximize log2FC_HepG2_ or log2FC_K562_, regardless of the other cell type (“Max1”), using the M0+1 model. The top 10 predicted sequences for each cell type were selected for inclusion in R2. The same procedure was applied using a clipped objective as above. 40 sequences total. 10 clipped and 10 unbounded designs per cell type (“Min1”) were also generated as above, this time minimizing log2FC_HepG2_ or log2FC_K562_, regardless of the other cell type. 40 sequences total.

### Ablations

Ablations were performed on the top 5 most specific enhancers measured in the R1 library. All possible single and pairwise double ablations of motifs were performed, unless a sequence only had 2 motifs, in which case the double ablation was not performed. Motif identification and coordinates were obtained via FIMO on the R1 library. To ablate a motif, the motif sequence was replaced with an equal-length subset of a dinucleotide-shuffling of the entire enhancer sequence. Dinucleotide shuffling was not performed on the motif sequence itself due to the high risk of motif-like elements emerging from the shuffling of a short motif sequence. 3 dinucleotide shuffles were performed per ablation. A 1 bp buffer upstream and downstream of the motif coordinates from FIMO were included in the ablation.

### Re-optimized R2 sequences

A masked version of Fast SeqProp was used to optimize all non-motif sequence while preserving motif sequence unchanged. The same enhancers and motif annotations used for the ablations were used here. 10 re-optimizations of each enhancer were generated. As a control, non-motif sequence was dinucleotide shuffled 5 times per enhancer.

### Target designs

4 target values were uniformly chosen in each cell type with a minimum |log2FC_H2K_| of 2 and maximum of the highest predicted specificity on R1 sequences using the M0+1 model. 100 sequences were generated per target, and the top 20 sequences with the closest predicted specificity to the target value were selected for inclusion in the R2 library. The Fast SeqProp objective was the same as for the Clipped designs. 160 sequences total.

### Shorter enhancer design

Enhancers with length <145bp were generated the same as for the Unbounded designs above, using 0-padding for the 200 bp model input size.

### Control sequence selection

The top 5 most specific sequences from R1 for each cell line were including in the R2 library. 100 sequences each were randomly selected from HepG2-targeted and K562-targeted R1 designs. 210 sequences total.

Additionally, the 2 sequences from R1 with lowest measured log2FC_HepG2_ + log2FC_K562_ were selected as negative controls. A dinucleotide shuffled variant of each of these sequences was also included in R2.

### Reduced library size estimations

To estimate the best enhancer for a given library (**Fig. 1F**), cell type, and library size n, n sequences were sampled with replacement 10,000 times, and the best log2FC_H2K_ for each bootstrap was then averaged together. For this analysis we used a subset of R0, filtered for sequences annotated with chromatin state = 5 (“Enhancer”, from a 25-state ChromHMM model^23^) to ensure the strongest comparison with synthetic enhancers; and treat the cell type of origin as the target cell type. For each of the 10,000 bootstraps, we also calculate which library (R0, R1, R1-DHS, R2) yields the best enhancer in each cell type, and report the percentage of bootstraps in which the best enhancer comes from each library.

### Ablation analysis

For each single and double ablation, 3 different dinucleotide shufflings of the target motif(s) were performed. The measurements for each ablation were averaged together, and the ablation score calculated on these averages. The deviation score was also calculated on the averaged measurements. For R2 Seq 976 and 1041, we estimated the double ablation score *in silico*. We finetuned the 9 individual M0+1 models on all R2 measurements (randomly split into 10 crossfolds) using the same hyperparameters used for R1 finetuning, then obtained ensemble predictions on 3 ablations per enhancer.

### Library synthesis and assembly

To test our synthetic library, we largely followed the same protocol used to generate the SHARPR-MPRA dataset^48^. Briefly, we filtered our designed enhancers to remove and replace any containing KpnI, XbaI, or SfiI recognition sites. Enhancers were placed next to their corresponding 3’UTR barcodes separated by KpnI and XbaI sites. We used 5 distinct barcodes per enhancer, which were generated using the dna-barcodes package (https://github.com/feldman4/dna-barcodes) (length 10, minimum edit distance = 2). Oligos containing enhancers and barcodes were ordered as a pool from Twist Biosciences. We amplified the oligo library with primers CY01 and CY02 (**Table S6**) to add Gibson overhangs; digested and gel purified the plasmid backbone (pMPRA1, Addgene #49349) with SfiI (NEB R0123) to remove its promoter and reporter sequence; then inserted the library cassette into the digested pMPRA1 backbone using the NEBuilder HiFi DNA Assembly kit (NEB E2621). This intermediate plasmid was purified with Kapa Pure Beads (Roche KK8002), transformed into NEB 10-beta electrocompetent cells (NEB C3020K), and extracted using the Qiagen Plasmid Maxi Kit (Qiagen 12162). We then insert a separate promoter and reporter cassette (minP, luciferase) excerpted from pMPRAdonor2 (Addgene #49353) in between the enhancer and barcode elements, by digesting both plasmids with KpnI (NEB R3142) and XbaI (NEB R0145), gel-purifying the appropriate fragments, dephosphorylating the intermediate plasmid fragment with Antarctic Phosphatase (NEB M0289) and ligating with T4 ligase (NEB M0202) for 16h. This ensures that barcodes will appear in RNA transcripts which can be mapped back to their corresponding enhancer. This final library plasmid was again purified, transformed, and Maxiprepped as above. The pooled library and 12 individual clones were sequence-verified via Sanger sequencing.

R1 and R2 libraries were synthesized via Twist and cloned as above. Given low variation observed among barcodes in R1, in R2 we reduced the number of barcodes per sequence from 5 to 2. To avoid PCR bias towards shorter sequences in R2, we used different primer sequence for 25 bp, 50 bp, 72 bp, and 145 bp enhancers. Each sublibrary was cloned separately into pMPRA1, then recombined according to the following proportions: 85% 145 bp, 5% 72 bp, 5% 50 bp, and 5% 25 bp. The second cloning step was performed on the recombined library.

### Library transfection

HepG2 cells were obtained from ATCC (HB-8065) and cultured in EMEM (ATCC 30-2003) + 10% FBS + 1% Pen/Strep at 37°C and 5% CO2. For each transfection, 600k (R1 libraries) or 1 million (R2) cells were seeded in 6-well plates or 6 cm plates respectively. 24 hours later, cells were transfected with Lipofectamine 3000 (Invitrogen L3000001) and 5 (R1) or 10 (R2) ug of library DNA following the manufacturer’s instructions. For R2, media was replaced 6 hours later. Cells were then grown for 48 hours after transfection. Transfection efficiency, estimated by transfecting a GFP plasmid in a parallel culture and performing flow cytometry, was ∼20%.

K562 cells were obtained from ATCC (CCL-243) and cultured in RPMI (Gibco 11875093) + 10% FBS + 1% Pen/Strep at 37°C and 5% CO2. For each transfection, 1.5 million K562 cells per replicate were electroporated with 10 ug DNA using the Neon Transfection System (Invitrogen MPK5000), transferred to 6-well plates (R1) or 6 cm plates (R2) with RPMI + FBS (no Pen/Strep), and incubated for 48 hours. Transfection efficiency, estimated by transfecting a GFP plasmid in a parallel culture and performing flow cytometry, was ∼90% (R1) and 96% (R2).

RNA was extracted 48h after transfection using the Monarch total RNA miniprep kit (NEB T2010S) and stored at −80C until further processing. Two independently-transfected replicates were performed per cell line, per DNA library.

### Library preparation and sequencing

mRNA was isolated from total RNA extracts using the magnetic mRNA Isolation Kit (NEB S1550S) following the manufacturer’s instructions. mRNA was then reverse transcribed with the Maxima H Minus Reverse Transcriptase (Thermo EP0753) using UMI-containing RT primer CY05 (**Table S6**), followed by digestion with RNase I (Thermo AM2294) and RNase H (NEB M0297) and purification with the DNA Clean & Concentrator - 5 (Zymo D4014), 7x binding buffer. To estimate the optimal number of PCR cycles for cDNA amplification, we first ran a small-scale qPCR for 30 cycles with 1-2 uL cDNA. qPCRs were ran with KAPA HiFi HotStart ReadyMix PCR Kit (Roche 07958935001), EvaGreen (Biotium 31000), and primers CY06/CY07 (fw) and Bri0xx (rev) containing P5 and P7 adapters and index sequences respectively (**Table S4**). The optimal cycle before the end of exponential amplification was determined from the amplification curves, and a new PCR reaction was run using this number with all remaining cDNA and scaling up all volumes accordingly. Amplification products were gel extracted using the Monarch® DNA Gel Extraction Kit (NEB T1020).

For DNA libraries, 500 ng plasmid library was amplified with HiFi HotStart ReadyMix PCR Kit and primers CY06/CY07 (fw) and CY05 (rev) for two cycles, then purified with the Zymo DNA Clean & Concentrator, 5x binding buffer. 50 ng of the resulting product were used in a small-scale qPCR with P5 and Bri0xx primers to determine the optimal amplification cycle as above. Finally, the remaining template was amplified via PCR using the optimal number of cycles determined via qPCR. Reaction products were gel purified as above.

For R1 libraries, we prepared 1 DNA replicate per enhancer library (R1-MPRA and R1-DHS). For R2 we prepared 2 DNA replicates. In all cases we prepared two RNA replicates per cell line, per enhancer library. Libraries were quantified using the KAPA Library Quantification Kit (Roche 7960140001) before mixing. Sequencing was performed in an Illumina NextSeq 550 with the following settings: Number of cycles for R1: >10 (enhancer barcode), R2: 10 (UMI), index 1: 8, index 2: 8, custom primers: Read 1: CY08, Read 2: Custom_read_2, Index 1: Custom_index_1, Index 2: CY009 (Custom_Index_2).

### Sequencing raw data processing

fastq file preprocessing, UMI deduplication, and barcode matching was performed using custom python scripts. UMI counts from barcodes corresponding to the same enhancers were pooled together. R1, R1-DHS, and R2 sequence measurements were processed with the DESeq2 package^49^, following the example of Zhao et al^10^. R0 sequence measurements were also re-processed with DESeq2 instead of total read count normalization prior to model retraining for R2 design. Enhancers were treated as genes, with cell type and replicate sublibraries treated as samples.

### Batch correction across design rounds

To improve our ability to compare measurements from different experiments, which covered a slightly different range in log2FC due to experimental variation, we performed a simple batch correction procedure as follows. Weighted linear regression was performed separately for each cell line on R2 measurements of R2 control sequences vs R1 measurements of the same sequences, using 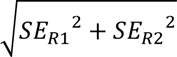 as the weights. For each cell type the following equation was obtained, where 𝑥 represents log2FC_HepG2_ or log2FC_K562_ and the subscript denotes library measurement: 𝑥_𝑅2_ = 𝑚_21_𝑥_𝑅1_ + 𝑏_21_. Regression slopes and intercepts (𝑚_10_, 𝑏_10_) were identically calculated for R1 vs R0.

Then, within each cell type batch-corrected R1 measurements were calculated as 𝑥_𝑅1∗_ = 𝑚_21_𝑥_𝑅1_ + 𝑏_21_; and batch-corrected R0 measurements were calculated as 𝑥_𝑅0∗_ = 𝑚_21_𝑚_10_𝑥_𝑅1_ + (𝑚_21_𝑏_10_ + 𝑏_21_). Standard errors were propagated appropriately, and log2FC_H2K_ scores recalculated on the batch-corrected measurements. All batch correction coefficients are reported in **Table S7.**

Finally, batch-corrected control sequence measurements were averaged together.

### Combinatorial indexing scMPRA library preparation

For HepG2, 2.5 million cells were electroporated with 11.5ug R1 library plasmid DNA in 250uL Resuspension Buffer R using the Neon Transfection System at 1230V, pulse width 20ms and 3 pulses total. 2 million cells were seeded in 2mL total media volume in a 6-well plate and fixed 24 hours later. For K562, 2.5 million cells were electroporated with 11.5ug R1 library plasmid DNA in 250uL Resuspension Buffer R using the Neon Transfection System at 1450V, pulse width 10ms and 3 pulses total. 2 million cells were seeded in 10mL total media volume in a T25 flask and fixed 24 hours later. We targeted a total of 10,000 transfected HepG2/K562 cells mixed at 1:1 ratio and performed 3-round combinatorial indexing using Parse Bio Evercode Mini v2 kit with modifications to the kit protocol for cDNA amplification steps to prepare an additional sequencing library enriching for MPRA amplicons. First, we spiked-in a MPRA amplicon targeting primer GB39 to initial cDNA amplification reaction at the final concentration of 400nM. We performed a second PCR, starting from 50ng of the cDNA from the initial amplification round, using GB42/GB43, with the following conditions: 0.02U/uL NEB Q5 HotStart Polymerase, 1x NEB Q5 Reaction Buffer, 300nM each primers, 20uM dNTPs, 1x Biotium EvaGreen dye; 98°C 20s - 67°C 10s - 72°C 30s. The cycling was stopped after the exponential phase, and the reaction products were double size-selected using Roche KAPA Pure beads at 0.8x-1.5x. Index PCR was performed according to the kit protocol alongside the whole transcriptome library. The final indexed libraries were quantified using KAPA Library Quantification Kit (Roche 7960140001) for pooling and sequenced at 2x150 cycles on NovaSeq X Plus by Novogene.

### scMPRA data processing

For both MPRA and transcriptome libraries, we initially performed anchored alignments to detect and filter the reads for correct amplicon structures using cutadapt^50^. For MPRA libraries, we extracted and concatenated only the barcode sequences from the reads and clustered them using starcode^51^ in order to remove PCR chimerism artifacts. We used STARsolo^52^ to align the reads to the human genome (hg38) and enhancer barcode references and to quantify the read counts. We filtered the cells based on a knee plot of per-cell transcriptome UMI counts. For the transcriptome data used for clustering and pseudobulk binning, we use normalized expression matrix following SCTransform scaling as implemented in Seurat^53^.

We observed that random barcodes were not an effective proxy for DNA copy numbers as indicated by: 1) the sparse, singleton-dominated per-plasmid expression levels (**Fig. S8E**); and 2) the poorer correlation of cell-type specific enhancer activities with the bulk data calculated using averages of random barcode-normalized single cell expressions compared to the correlation calculated using pseudobulk (**Fig. S8F**). Thus, we used pseudobulk quantifications in our downstream analysis.

## Data and code availability

Raw sequencing data along with unnormalized read counts are available in the Gene Expression Omnibus repository under accession code GSE269036 for R1-MPRA, R1-DHS, and R2; and accession code GSE269037 for the scMPRA experiment. Code used to train models, design sequences, analyze data, and produce figures are made available at https://github.com/christopheryin/iterative_synthetic_enhancer_design, along with processed data files.

