## Supplementary material for "Iterative deep learning-design of human enhancers exploits condensed sequence grammar to achieve cell type-specificity": Figures S1-11

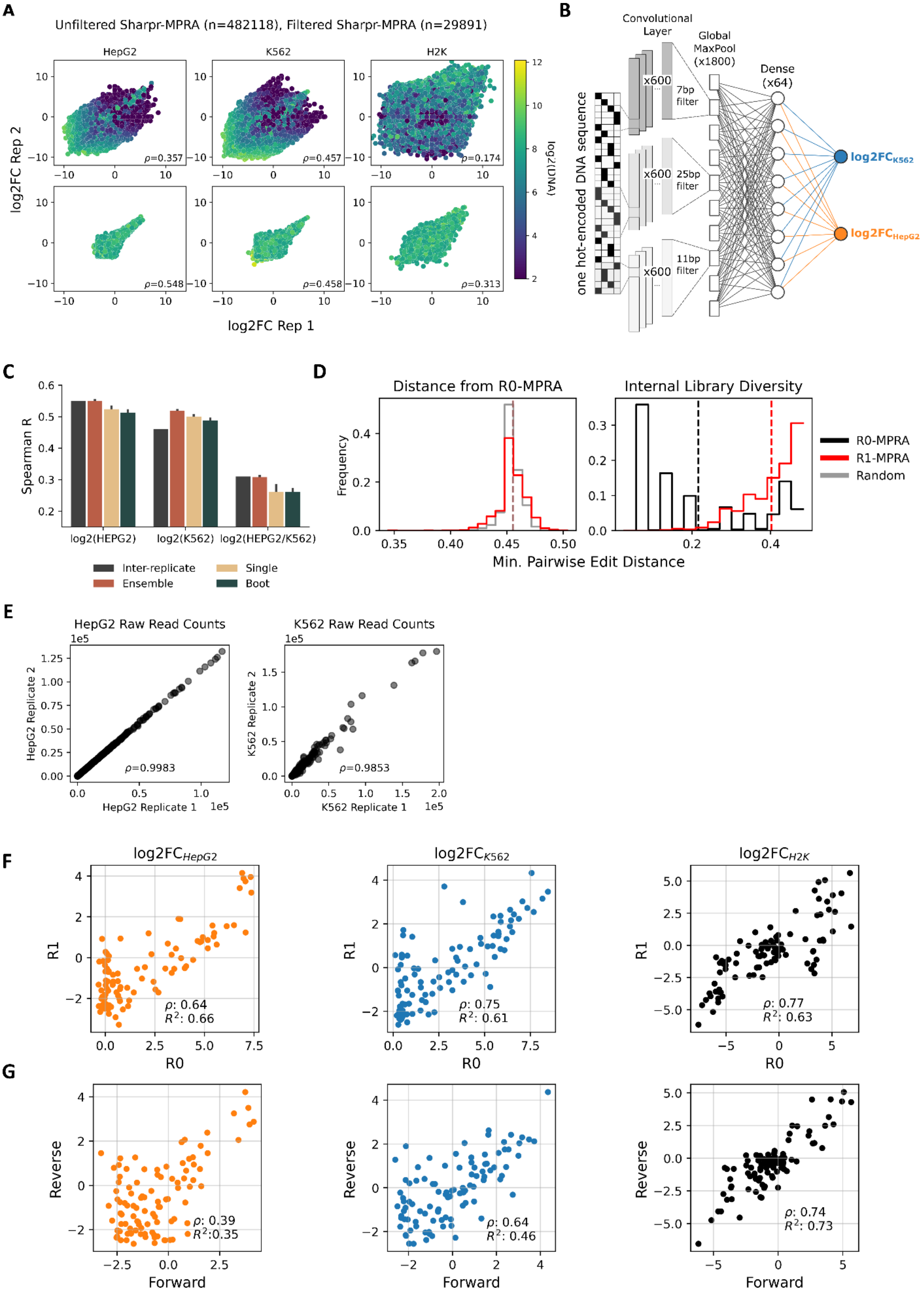
A
Unfiltered Sharpr-MPRA (n=482118), Filtered Sharpr-MPRA (n=29891)
HepG2
K562
H2K
log2FC Rep 2
log2FC Rep 1
log2(DNA)
ρ=0.357
ρ=0.457
ρ=0.174
ρ=0.548
ρ=0.458
ρ=0.313
B
Convolutional Layer
Global MaxPool (x1800)
Dense (x64)
one hot-encoded DNA sequence
x600
7bp filter
25bp filter
11bp filter
log2FCK562
log2FCHepG2
C
Spearman R
log2(HEPG2)
log2(K562)
log2(HEPG2/K562)
Inter-replicate
Ensemble
Single
Boot
D
Distance from R0-MPRA
Internal Library Diversity
Frequency
Min. Pairwise Edit Distance
R0-MPRA
R1-MPRA
Random
E
HepG2 Raw Read Counts
K562 Raw Read Counts
HepG2 Replicate 2
HepG2 Replicate 1
K562 Replicate 2
K562 Replicate 1
ρ=0.9983
ρ=0.9853
F
log2FCHepG2
log2FCK562
log2FCH2K
R1
R0
ρ: 0.64
R²: 0.66
ρ: 0.75
R²: 0.61
ρ: 0.77
R²: 0.63
G
Reverse
Forward
ρ: 0.39
R²:0.35
ρ: 0.64
R²: 0.46
ρ: 0.74
R²: 0.73

**Fig. S1 R0-MPRA processing and R1-MPRA generation. | A**, Inter-replicate correlation (Spearman, indicated with the letter "ρ") for $log2FC_{HepG2}$, $log2FC_{K562}$, and $log2FC_{H2K}$ in unfiltered Sharpr-MPRA (top row) vs. filtered Sharpr-MPRA (bottom row, R0), using total read count normalized measurements. Points are colored by log2(DNA count) . **B**, Architecture of multitask CNN model trained on R0-MPRA data and used to design R1-MPRA sequences. **C,** Prediction-measurement correlation and inter-replication correlation compared on $log2FC_{HepG2}$, $log2FC_{K562}$, and $log2FC_{H2K}$ held-out test set measurements for Single, Boot, and Ensemble model types trained on R0-MPRA sequences (10 models each). **D**, Left: Minimum pairwise edit distance between each R1-MPRA sequence and all R0-MPRA sequences, as well as between all R0-MPRA sequences and a library of 1000 sequences randomly generated from R0-MPRA nucleotide frequencies. Right: Minimum pairwise edit distance between all R1-MPRA sequences compared to within all R0-MPRA sequences. Means plotted as vertical dashed lines. **E,** Inter-replicate correlation for mRNA raw read counts in each cell type for R1-MPRA library. **F,** Correlation between R0-MPRA (reprocessed from measurements in Ernst *et al.*[23])and R1-MPRA (our) measurements of the 100 control sequences shared between libraries. **G**, Correlation between measurements of forward and reverse complement of 100 control sequences in R1-MPRA.

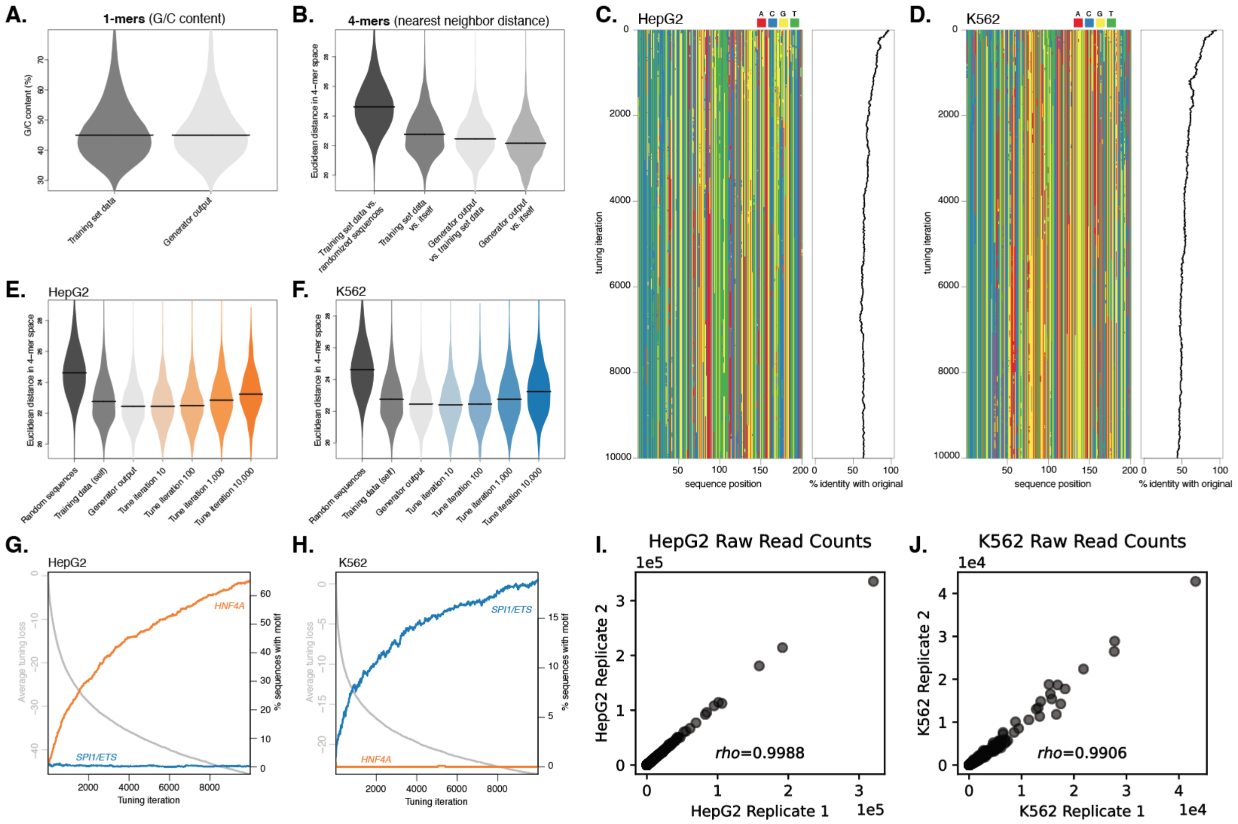


**Fig. S2 R1-DHS generation. | A-B**, Characteristics of endogenous accessible sequence elements used for training the GAN vs. synthetic GAN-generated sequences w.r.t. G/C content percentage (**A**) and general 4-mer sequence content (**B**), the latter reporting Euclidean distances between indicated pairs of sequence sets. **C-D**, Example sequence tuning of a single GAN-generated sequence towards a HepG2 (**C**) and K562 (**D**) cellular context. Colormap shows the tuning process, starting from the same generated sequence, and slowly converging to cell type-specific sequences, while retaining a large fraction of the original sequence identity. **E-F,** Sequence characteristics during tuning iterations for HepG2 (**E**) and K562 (**F**), as shown by Euclidean distance in 4-mer space for indicated sequence sets, training data. **G-H,** Average tuning loss across tuning iterations versus percentage of sequences containing select transcription factor motifs. Shown are HNF4A (orange) and SPI1/ETS (blue), the former being enriched in HepG2 cells (**G**) and the latter in K562 cells (**H**). **I-J,** Inter-replicate correlation for mRNA raw read counts for the R1-DHS library in HepG2 (**I**) and K562 (**J**).

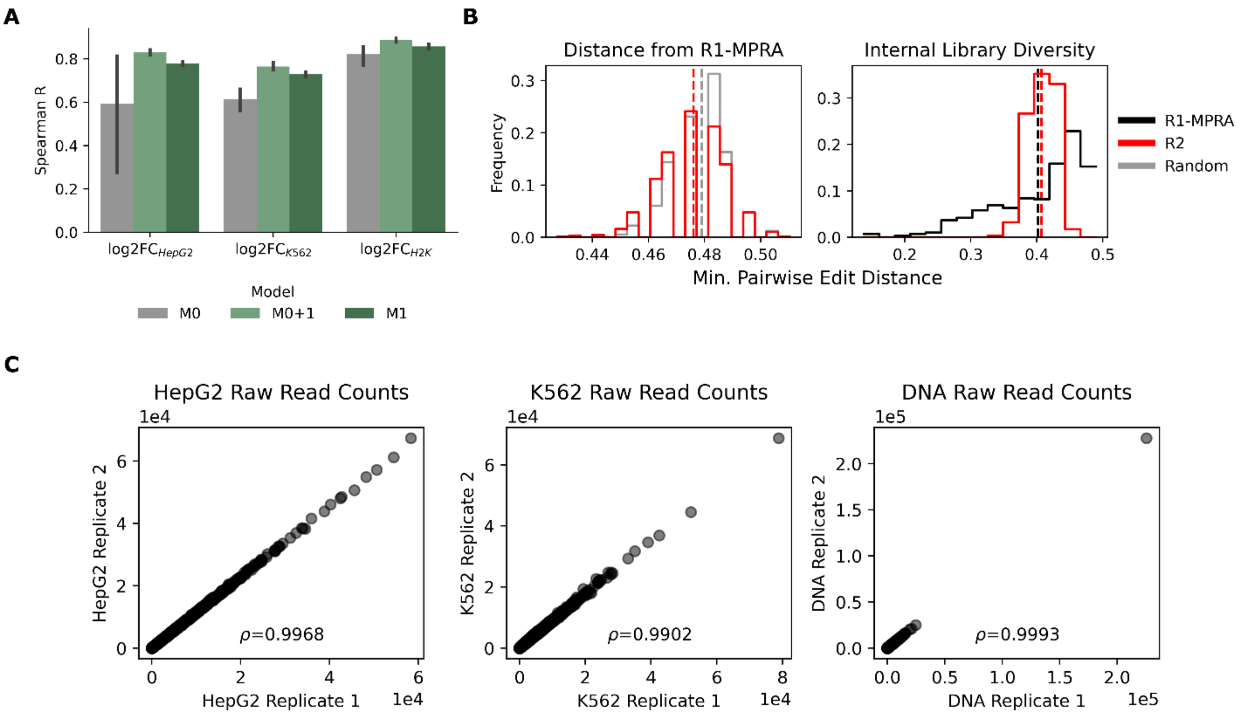


**Fig. S3 R2 generation. | A**, Prediction performance on held-out test set of R1-MPRA data for models trained on R0 data only ("M0"), M0 models finetuned on R1 data ("M0+1"), and models trained only on R1 data ("M1"). **B,** Minimum pairwise edit distance between each R2 sequence and all R1-MPRA sequences, as well as between all R1-MPRA sequences and a library of 1000 sequences randomly generated from R1-MPRA nucleotide frequencies (left). Minimum pairwise edit distance between all R2 sequences compared to within all R1-MPRA sequences (right). Means plotted as dashed lines. **C,** Inter-replicate correlation (Spearman, indicated with the letter "ρ") for raw read counts in each cell type for R2 library.

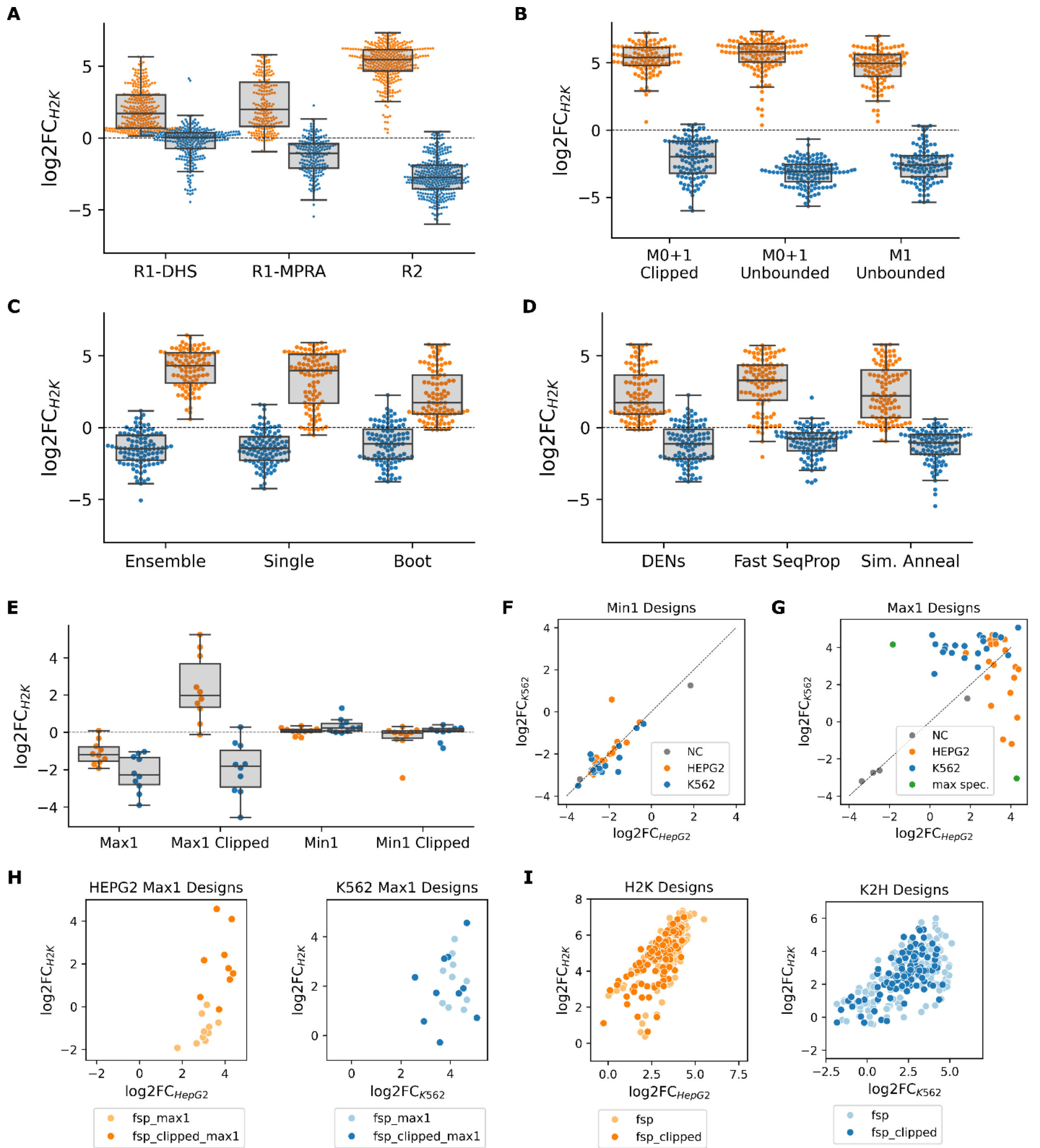


**Fig. S4 Comparing design practices. | A-D**, Comparing enhancer performance between sublibraries within R1 and R2. Each marker corresponds to an enhancer, colored by target cell type. **A,** Comparing enhancers designed to maximize specificity across R1-MPRA, R1-DHS, and R2. **B**, Comparing R2 designs. M0+1 models designed with clipped vs. unbounded objective (left), M0+1 models vs M1 models (right). **C**, Comparing model types in R1 for DEN-generated sequences (the only design method shared across all model types). **D**, Comparing design methods in R1 for Boot-generated sequences (the only model type with comparable numbers of sequences generated from each design method). **E**, Boxswarm plots of log2FC$_{H2K}$ for Max1 and Min1 designs. Max1 designs in K562 had moderate specificity (median log2FC$_{H2K}$ = -2.28, -1.81 for unbounded and clipped objectives); whereas Max1 designs in HepG2 had lower on-target specificity with the unbounded objective (-1.20) compared to moderate specificity with the clipped (1.98) **F**, Scatterplot of $log2FC_{K562}$ vs $log2FC_{HepG2}$ for Min1 designs (includes clipped and unbounded objectives), as well as 4 negative control (NC) sequences. Line of unity shown. **G**, Scatterplot of $log2FC_{K562}$ vs $log2FC_{HepG2}$ for Max1 designs (includes clipped and unbounded objectives), as well as 4 negative control (NC) sequences. Additionally, the most specific 145bp synthetic enhancer in each cell type is plotted in green (max spec.). Line of unity shown. **H**, Specificity vs target cell type strength for Max1 designs, compared to **I**, Specificity vs target cell type strength for R2 enhancers designed to maximize specificity, same as Fig. 1E.

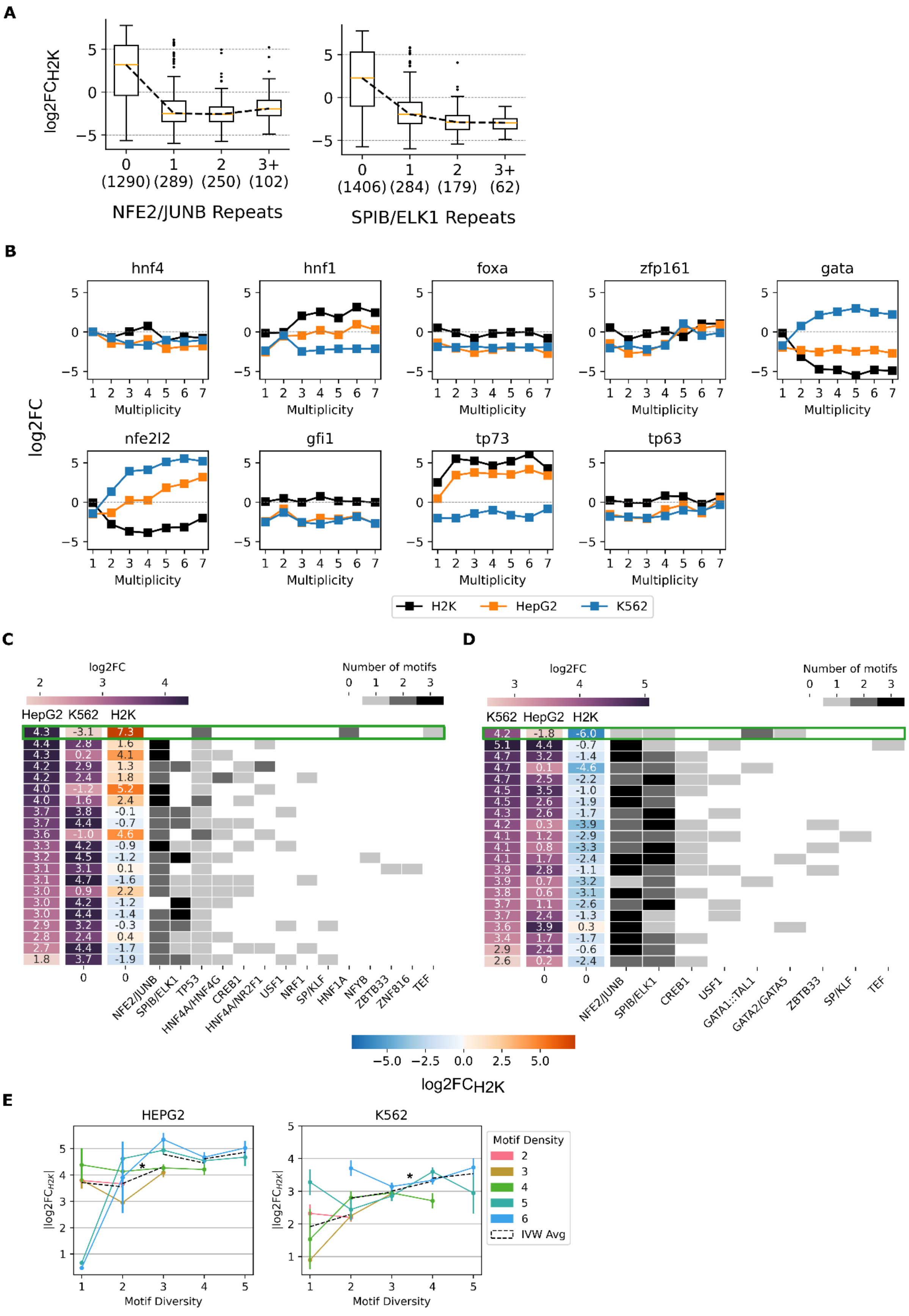
A
log2FCH2K
0
1
2
3+
(1290) (289) (250) (102)
NFE2/JUNB Repeats
(1406) (284) (179) (62)
SPIB/ELK1 Repeats
B
hnf4
hnf1
foxa
zfp161
gata
nfe2l2
gfi1
tp73
tp63
log2FC
Multiplicity
H2K
HepG2
K562
C
log2FC
Number of motifs
HepG2 K562 H2K
NFE2/JUNB
SPIB/ELK1
TP53
HNF4A/HNF4G
CREB1
HNF4A/NR2F1
USF1
NRF1
SP/KLF
HNF1A
NFYB
ZBTB33
ZNF816
TEF
D
log2FC
Number of motifs
K562 HepG2 H2K
NFE2/JUNB
SPIB/ELK1
CREB1
USF1
GATA1::TAL1
GATA2/GATA5
ZBTB33
SP/KLF
TEF
log2FCH2K
E
HEPG2
K562
|log2FCH2K|
Motif Diversity
Motif Density
2
3
4
5
6
IVW Avg

**Fig. S5 Additional motif analysis. | A**, log2FC$_{H2K}$ vs multiplicity for motifs not featured in main text. **B**, log2FC$_{H2K}$, log2FC$_{HepG2}$, and log2FC$_{K562}$, vs multiplicity for manual homotypic enhancers measured in R1. Each point corresponds to a single sequence measurement. **C,D** Motif content of HepG2 (**D**) and K562 (**E**) Max1 designs plotted alongside log2FC$_{H2K}$, log2FC$_{HepG2}$, and log2FC$_{K562}$. The most specific 145bp synthetic enhancer from R2 is plotted at the top of each heatmap for reference (green rectangle), otherwise sequences sorted by descending strength in target cell type. NFE2/JUNB, SPIB/ELK1, CREB1, and USF1 motifs prominently deployed in Max1 designs targeted to both cell types, indicating some level of non-specific activity. **E**, Mean log2FC$_{H2K}$ vs motif diversity (number of unique motif types in a sequence) plotted for R2 enhancers stratified by motif density. Best fit lines shown for all stepwise increases in motif diversity, within each strata. Inverse variance-weighted average slopes ("IVW Avg") across strata plotted as dashed black lines. In HepG2 designs a significant positive IVW Avg slope is observed when increasing motif diversity from 2 to 3 (m = 0.645+-0.178, p = 3e-4, denoted by asterisk). In K562 designs a significant positive IVW Avg slope is observed when increasing motif diversity from 3 to 4 (m = 0.297+-0.117, p = 1e-2, denoted by asterisk), and when regressing across all stepwise increases in motif diversity (m = 0.244+-0.058, p = 2.7e-5, line not shown).

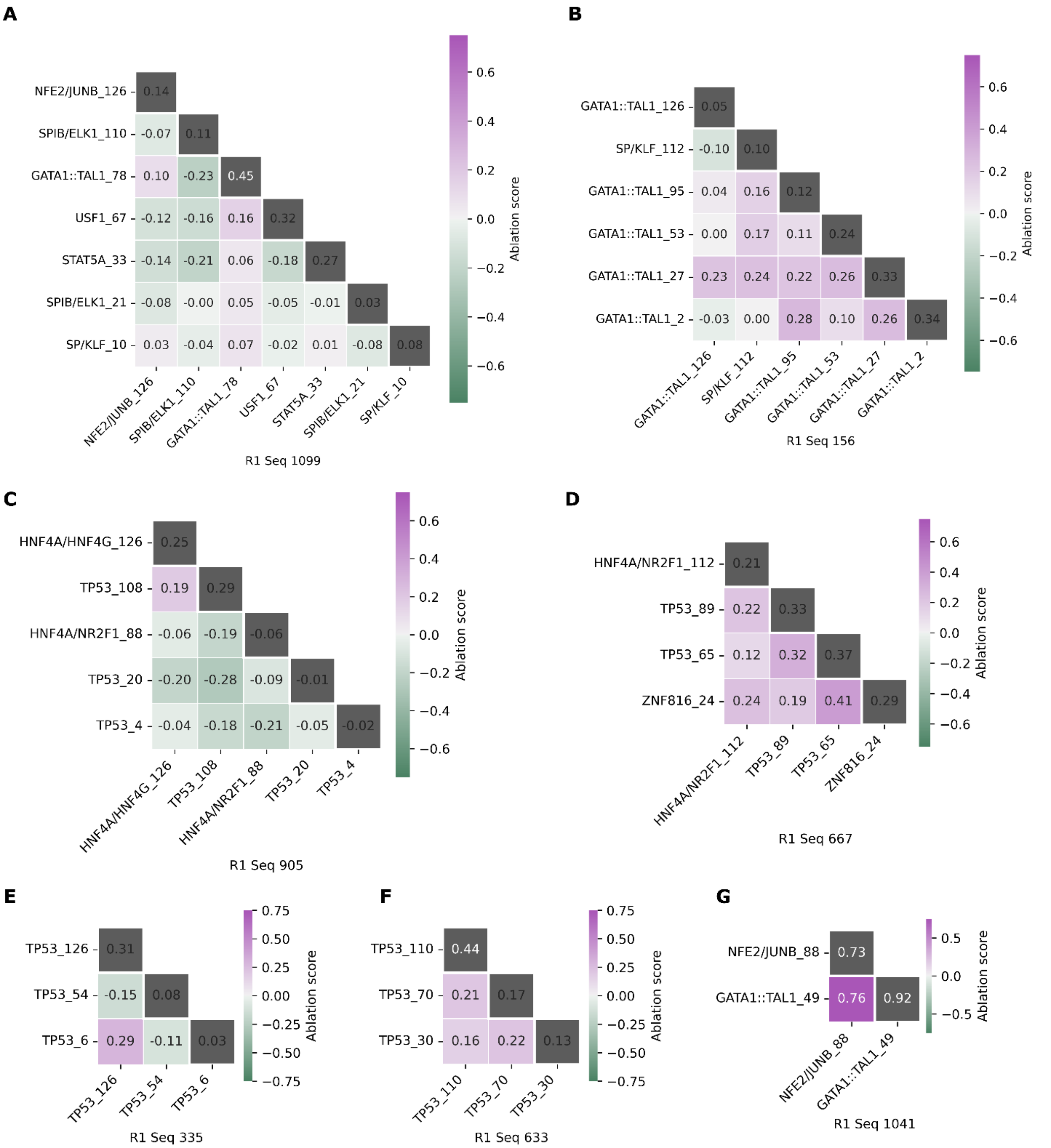


**Fig. S6 Additional double ablation deviation score heatmaps for sequences in Fig. 3. | A**, R1 Seq 1099. **B**, R1 Seq 156. **C**, R1 Seq 905. **D**, R1 Seq 667. **E**, R1 Seq 335. **F**, R1 Seq 633. **G**, R1 Seq 1041. See also **Fig. 3F-H**.

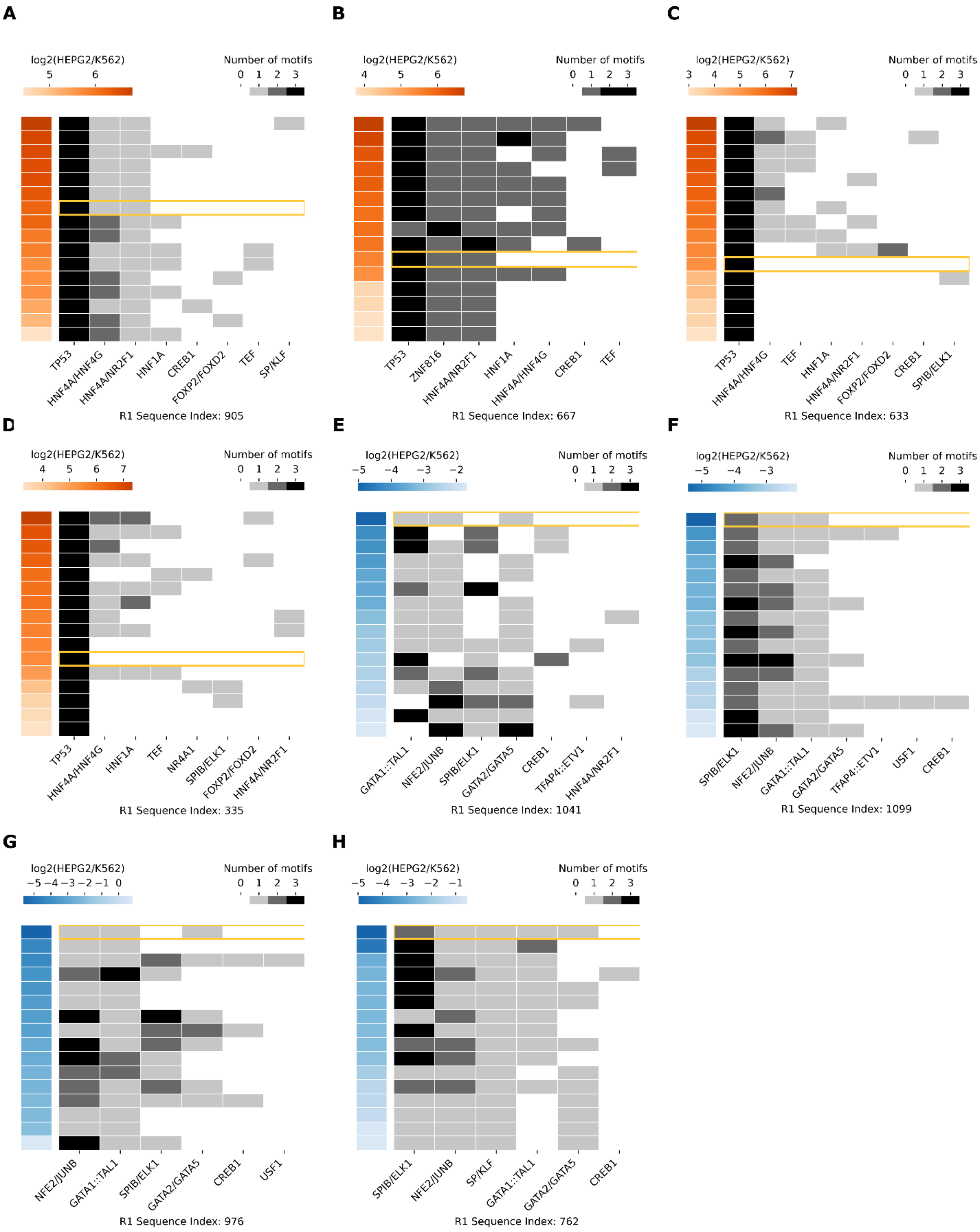


**Fig. S7 Additional motif content of non-motif redesigned enhancers in Fig. 3. | A**, R1 Seq905. **B**, R1 Seq 667. **C**, R1 Seq 633. **D**, R1 Seq 335. **E**, R1 Seq 1041. **F**, R1 Seq 1099. **G**, R1 Seq 976. **H**, R1 Seq 762. See also **Fig. 3I-K**.

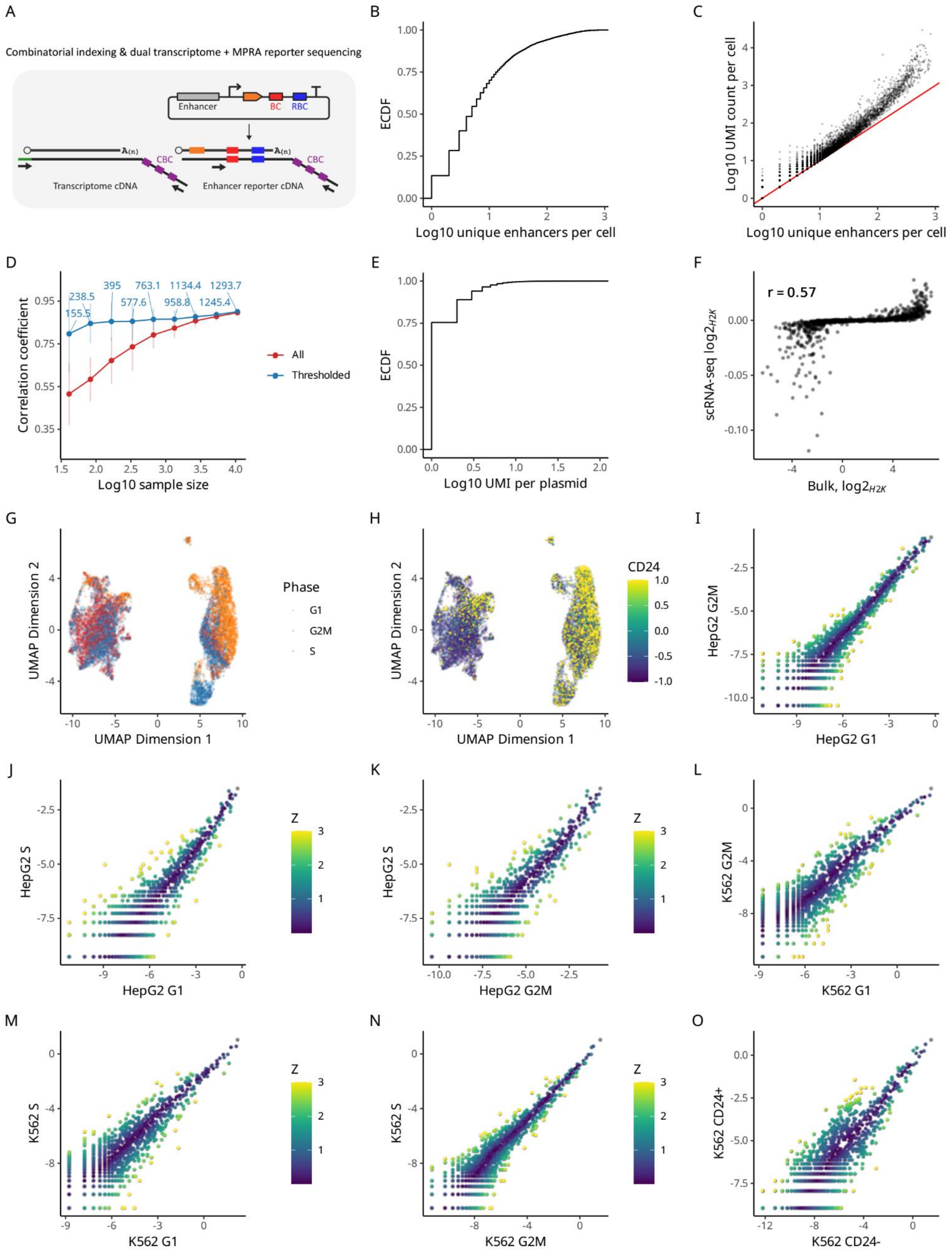

A
Combinatorial indexing & dual transcriptome + MPRA reporter sequencing
Enhancer
BC
RBC
A(n)
CBC
Transcriptome cDNA
Enhancer reporter cDNA
B
ECDF
Log10 unique enhancers per cell
C
Log10 UMI count per cell
Log10 unique enhancers per cell
D
155.5
238.5
395
577.6
763.1
958.8
1134.4
1245.4
1293.7
Correlation coefficient
Log10 sample size
All
Thresholded
E
ECDF
Log10 UMI per plasmid
F
r = 0.57
scRNA-seq log2$_{H2K}$
Bulk, log2$_{H2K}$
G
UMAP Dimension 2
UMAP Dimension 1
Phase
G1
G2M
S
H
CD24
UMAP Dimension 2
UMAP Dimension 1
I
HepG2 G2M
HepG2 G1
Z
J
HepG2 S
HepG2 G1
Z
K
HepG2 S
HepG2 G2M
Z
L
K562 G2M
K562 G1
Z
M
K562 S
K562 G1
Z
N
K562 S
K562 G2M
Z
O
K562 CD24+
K562 CD24-
Z

**Fig. S8 scRNA quality analysis. A,** A schematic illustrating scMPRA via combinatorial indexing. Cells are transfected with a modified enhancer reporter library that has random barcodes to tag individual plasmid molecules. Single-cell RNA-seq is carried out via combinatorial indexing, with separate PCRs to amplify both the transcriptome and reporter library cDNA amplicons. BC: enhancer barcodes; RBC: random barcodes; CBC: cell barcodes. **B,** Empirical CDF of the distribution of the numbers of unique enhancer sequences detected per cell. **C,** A scatter plot illustrating total UMI counts per cell versus the numbers of unique enhancer sequences per cell. **D,** Subsampling analysis of bulk vs. pseudobulk correlations. Cells used for pseudobulk aggregation are subsampled at various cell numbers as shown on the X-axis. Mean and standard deviation of bulk vs. pseudobulk correlations from n=100 samples for each subsample sizes are plotted as dots and error bars. In blue are the correlations calculated with enhancers passing minimum reporter expression level threshold (>4 UMI / cell), with the text label indicating the average number of enhancers that were available to calculate the correlations for each subsample size at this threshold. **E,** Empirical CDF of the distribution of the numbers of UMIs expressed per unique plasmid molecule. **F,** A scatter plot comparing cell-type specific activity of synthetic enhancers obtained by bulk vs. single-cell analysis, using mean RBC-normalized single cell enhancer expression. **G,** UMAP projection of the single cell transcriptomes, colored by cell cycle phases. **H,** UMAP projection of the single cell transcriptomes, colored by CD24 expression, a differentiation marker in K562 cells. **I-O,** Pairwise scatter plots comparing enhancer expression between substates (cell cycle phases and K562 CD24+/- status) within each cell types. Colored by z-scaled log2 fold differences between the aggregated pseudobulk counts on X and Y axes.

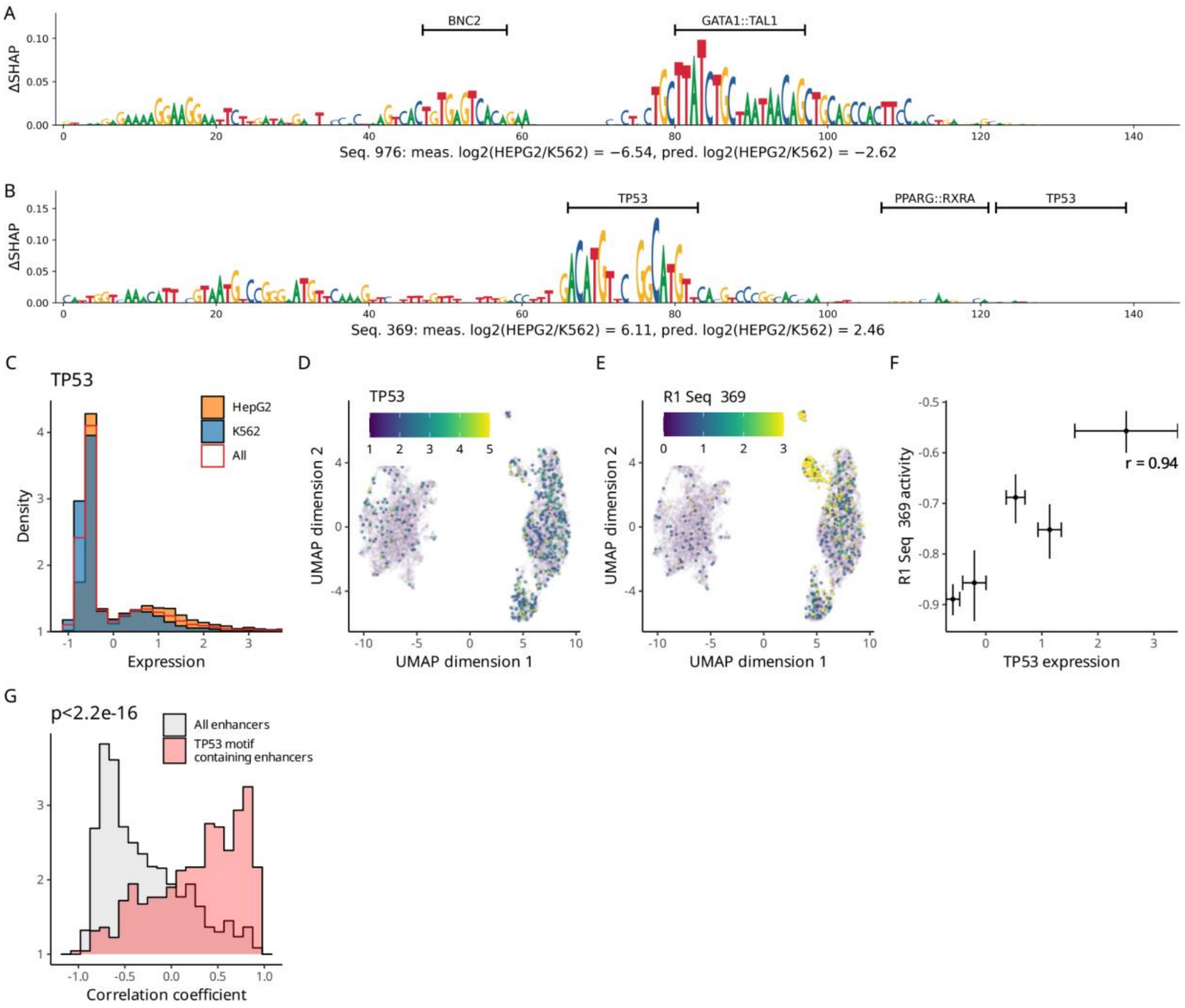


**Fig. S9 Single cell level enhancer activity. A,** A sequence logo for R1 Seq 976 with nucleotide height corresponding to the difference between $log2FC_{K562}$ SHAP value and $log2FC_{HepG2}$ SHAP value, according to the M0+1 ensemble (**Methods**). **B,** A sequence logo for R1 Seq 368 with nucleotide height corresponding to the difference between $log2FC_{HepG2}$ SHAP value and $log2FC_{K562}$ SHAP value, according to the M0+1 ensemble (**Methods**). **C,** Expression levels of TP53 transcription factor across: all cells in red; cells in HepG2 cluster in orange; cells in K562 cluster in blue. **D,** Expression levels of TP53 transcription factors across single cells atop the UMAP projection of the transcriptomes. **E,** Activities of an enhancer #369, containing a TP53 transcription factor motif, across single cells atop the UMAP projection of the transcriptomes. **F,** Pseudobulk correlation of TP53 transcription factor expression levels and enhancer #369. The pseudobulks are binned by TP53 expression levels. Vertical error bars represent bootstrap standard errors of enhancer activities. Horizontal error bars represent standard deviation of the transcription factor expression levels across the cells in each pseudobulk bin. **G,** Distribution of correlations between TP53 expression versus all enhancers across pseudobulk of single cells binned by TP53 expression values. In red are pairings where the enhancer contains the DNA sequence motif for TP53. p value from Kolmogorov-Smirnov test.

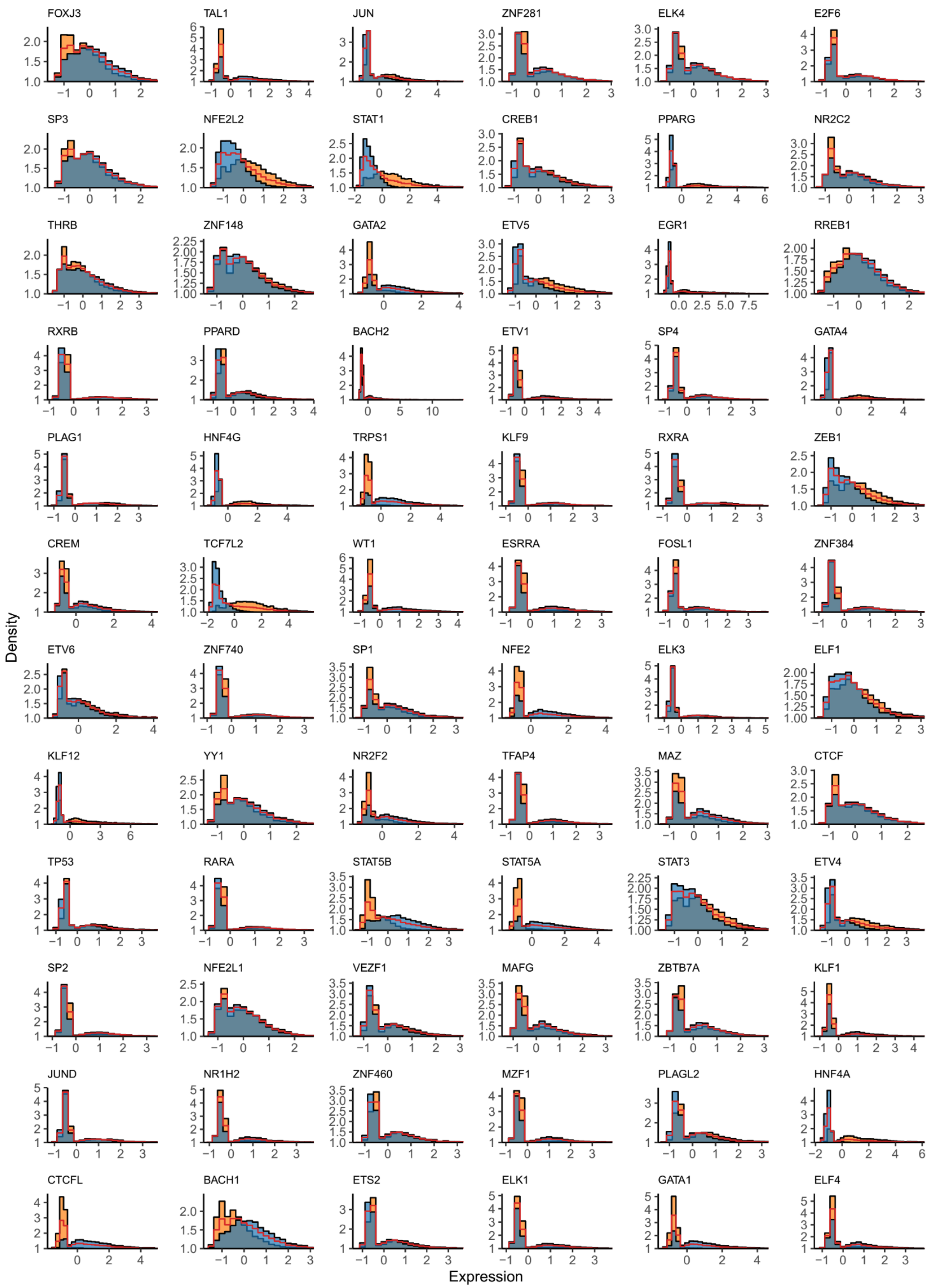


**Fig. S10 Expression levels of TF transcription factors.** Expression across all cells in red; cells in HepG2 cluster in orange; cells in K562 cluster in blue.

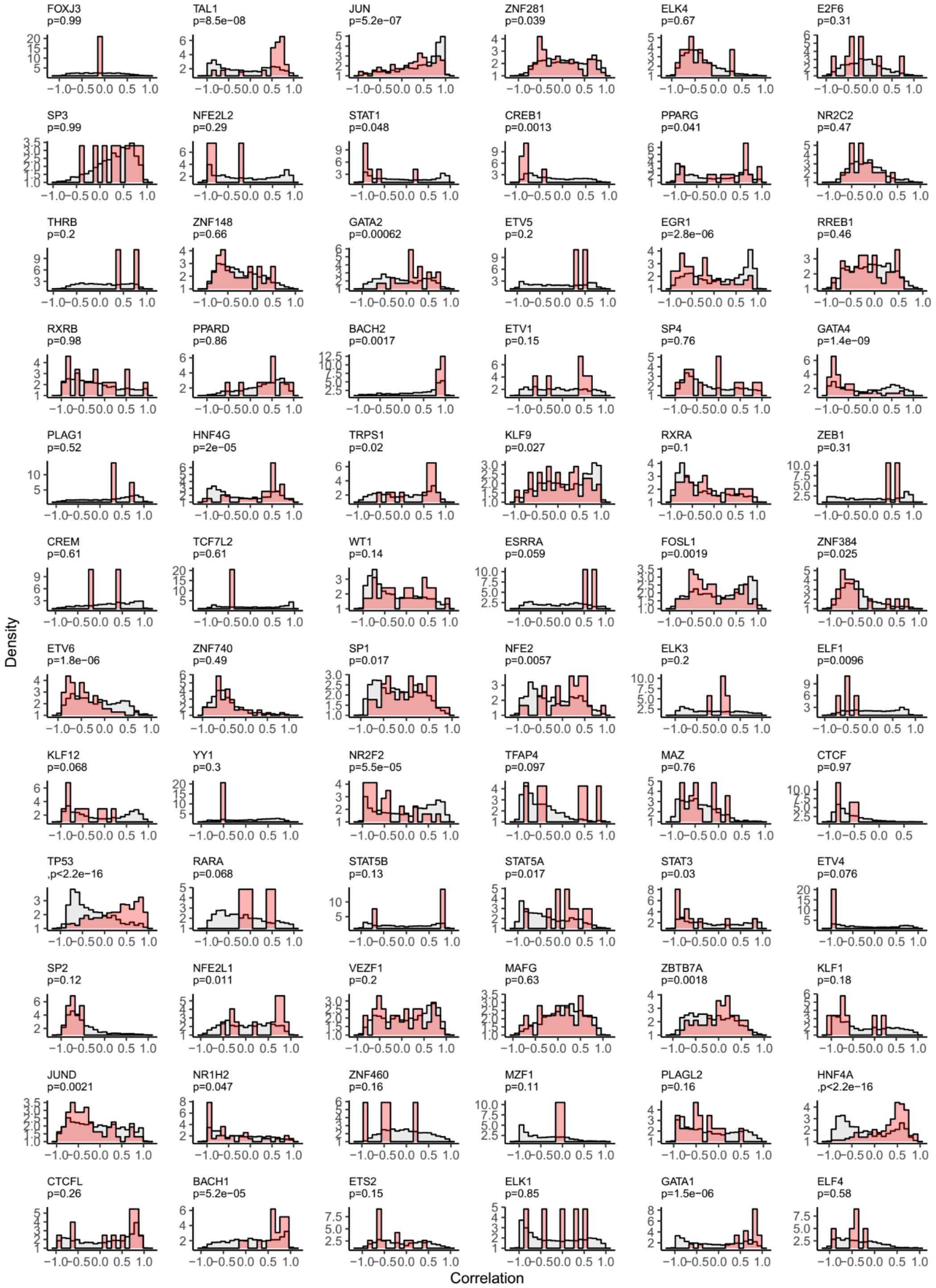


**Fig. S11 Correlation between TF expression and enhancer activity.** Distribution of correlations between TF expression versus all enhancers across pseudobulk of single cells binned by TF expression values. In red are pairings where the enhancer contains the DNA sequence motif for each TF. p values from two-sided Kolmogorov-Smirnov test. Benjamini-Hochberg estimated FDR<0.05 at p<0.041.
