## Supplementary material for "Iterative deep learning-design of human enhancers exploits condensed sequence grammar to achieve cell type-specificity": Tables S1-7

| | Fast SeqProp | Sim. Anneal | DEN | Other |
|---|---|---|---|---|
| Single | 28 | 18 | 200 | - |
| Boot | 198 | 198 | 198 | - |
| Ensemble | - | - | 198 | - |
| Control | - | - | - | 200 |
| Motif repeat | - | - | - | 62 |

**Table S1 – R1-MPRA Library Composition |** Numbers in cells indicate the total number of sequences in R1-MPRA generated using a given model type (row) and design type (column).

| Model type | Design type | Design objective | Count |
|---|---|---|---|
| M1 | Fast SeqProp | Unbounded H2K | 210 |
| M0+1 | Fast SeqProp | Unbounded H2K | 258 |
| | | Clipped H2K | 220 |
| | | Target H2K | 160 |
| | | Unbounded Max1 | 20 |
| | | Unbounded Min1 | 20 |
| | | Clipped Max1 | 20 |
| | | Clipped Min1 | 19 |
| | Masked Fast SeqProp (nonmotif reoptimization) | Unbounded H2K | 100 |
| | Reduced length Fast Seqprop | Unbounded H2K | 110 |
| None | Dinucleotide shuffling | Nonmotif reoptimization | 49 |
| | | Motif ablation | 331 |
| Control | Negative control | - | 4 |
| | Random | - | 198 |
| | Top enhancer | - | 10 |

**Table S2 – R2 Library Composition |** Breakdown of R2 library composition.

| CISBP2.0 Motif Name | Sequence (5’-3’) | Putative Role |
|---|---|---|
| Hnf4 | AGGTTCAAAGGTCA | HepG2 Enhancer |
| Hnf1 | GGTAATTATTAACC | HepG2 Enhancer |
| Foxa | TGTTTACTTAGG | HepG2 Enhancer |
| Zfp161 | TGGCGCGCGCGCCTGA | K562 Enhancer |
| Gata | CTGGTGGGGACAGATAAG | K562 Enhancer |
| Nfe2l2 | ATGACTCAGCA | K562 Enhancer |
| Gfi1 | AAATCACAGC | K562 Repressor |
| Tp73 | ACATGTC | HepG2 Enhancer |
| Tp63 | ACATGCCCGGGCATG | HepG2 Enhancer |

**Table S3 – Motif sequences used in hand-crafted motif repeat enhancers**

| Generator Layers: | Dimension (BS x W x H x C) |
|---|---|
| Input: noise / seed | N x 100 |
| Linear | N x 1600 |
| ReLU | N x 1600 |
| BatchNorm | N x 1600 |
| Transposed Conv (num_filters = 640, filter size = 10x1, stride = 10) | (N x 200 x 1 x 640) |
| ReLU | (N x 200 x 1 x 640) |
| BatchNorm | (N x 200 x 1 x 640) |
| Transposed Conv (num_filters = 1, filter size = 15x4, stride = 1) | (N x 200 x 4 x 1) |
| Softmax | (N x 200 x 4 x 1) |

| Discriminator Layers: | Dimension (BS x W x H x C) |
|---|---|
| Input: one hot sequence | (N x 200 x 4 x 1) |
| Conv (num_filters = 640, filter size = 15x4, stride = 1) | (N x 200 x 1 x 640) |
| Spectral Norm | (N x 200 x 1 x 640) |
| Max Pool (kernel size = 10x1) | (N x 20 x 1 x 640) |
| Leaky ReLU (slope=0.1) | (N x 20 x 1 x 640) |
| Conv (num_filters = 320, filter size = 11x1, stride = 1) | (N x 20 x 1 x 320) |
| Spectral Norm | (N x 20 x 1 x 320) |
| Max Pool (kernel size = 20x1) | (N x 1 x 1 x 320) |
| Leaky ReLU (slope=0.1) | (N x 1 x 1 x 320) |
| Linear (200) | N x 200 |
| Spectral Norm | N x 200 |
| Leaky ReLU (slope=0.1) | N x 200 |
| Linear(1) | N x 1 |

**Table S4 ー Model architecture of Generative Adversarial Network (GAN)** | (N = batch size, dimension format: batch size x width x height x channels)

| **Layers** | **Dimension (BS x W x H x C)** |
| --- | --- |
| (input) | (N x 200 x 4 x 1) |
| Conv (num filters = 32, filter size = 15x4, stride = 1) | (N x 200 x 1 x 32) |
| ReLU | (N x 200 x 1 x 32) |
| Dropout (dropout rate = 0.5) | (N x 200 x 1 x 32) |
| Max Pool (kernel size = 200) | (N x 1 x 1 x 32) |
| Linear (50) | N x 50 |
| ReLU | N x 50 |
| Dropout (dropout rate = 0.5) | N x 50 |
| Batch Normalization | N x 50 |
| Linear (3) | N x 3 |
| Softmax | N x 3 |

**Table S5 — Model architecture of Classification model** | (N = batch size, dimension format: batch size x width x height x channels)

| Primer Name | Primer Description | Primer Sequence (5'-3') |
|---|---|---|
| CY01 | Add Gibson flank to 5' of R1-MPRA and R2 145bp enhancer, for overlap with pMPRA1 backbone | gaacatttctctGGCCTAACTGGCCGCTTCACTG |
| CY02 | Add Gibson flank to 3' end of R1-MPRA enhancer, for overlap with pMPRA1 backbone | cccgactagcttggccgccgtGGCCCCGCTCCTGTATAGCTG |
| CY03 | Add Gibson flank to 5' of R1-DHS enhancer, , for overlap with pMPRA1 backbone | gaacatttctctGGCCTAACTGGCCCTTCGCTG |
| CY04 | Add Gibson flank to 3' end of R1-DHS enhancer, for overlap with pMPRA1 backbone | cccgactagcttggccgCCGTGGCCTCAGTTCACCGCGTC |
| CY05 | Reverse transcriptase primer (adds UMI to mRNA transcript) | AAGCAGTGGTATCAACGCAGAGTACATGGGNNNNNNNNNNccaaactcatcaatgtatcttatcatgt |
| CY06 | qPCR forward R1-MPRA enhancer | AATGATACGGCGACCACCGAGATCTACACGGCTCTGAtctcattaaggccaagaagggc |
| CY07 | qPCR forward R1-DHS | AATGATACGGCGACCACCGAGATCTACACAGGCGAAGtctcattaaggccaagaagggc |
| CY08 | Custom read1 | ggcggcaagatcgccgtgtaataattCTAGA |
| CY09 | Custom Index 2 | tgccgcccttcttggccttaatgaga |
| CY10 | Add Gibson flank to 5' of R2 25bp enhancer, for overlap with pMPRA1 backbone | gaacatttctctGGCCTACCTATGCCCACGTCCC |
| CY11 | Add Gibson flank to 3' of R2 25bp enhancer, for overlap with pMPRA1 backbone | cccgactagcttggccgccgGTGATACGTGTGTCTGGC |
| CY12 | Add Gibson flank to 5' of R2 50bp enhancer, for overlap with pMPRA1 backbone | gaacatttctctGGCCTATCGGAATCGGTAACGGC |
| CY13 | Add Gibson flank to 3' of R2 50bp enhancer, for overlap with pMPRA1 backbone | cccgactagcttggccgccgGTACACCACTGTCCACTG |
| CY14 | Add Gibson flank to 5' of R2 72bp enhancer, for overlap with pMPRA1 backbone | gaacatttctctGGCCTATTCCATCCGCCTGACC |
| CY15 | Add Gibson flank to 3' of R2 72bp enhancer, for overlap with pMPRA1 backbone | cccgactagcttggccgccgGTCTGTGAGCATCGACC |
| SC_01264 | Add Gibson flank to 3' end of R2 145bp enhancer, for overlap with pMPRA1 backbone | cccgactagcttggccgccgATCTACCTGGTCCGGCA |
| Custom_read_2 | | AAGCAGTGGTATCAACGCAGAGTACATGGG |
| Custom_index_1 | | CCCATGTACTCTGCGTTGATACCACTGCTT |
| GB_39 | | GGCAAGATCGCCGTGTAATAATTCTAGA |
| GB_42 | | TCGTCGGCAGCGTCAGATGTGTATAAGAGACAGGGCAAGATCGCCGTGTAATAATTCTAGA |
| GB_43 | | GTGACTGGAGTTCAGACGTGTGCTCTTCCGATCT |
| GB_29 | | cggccaagctagtcggggcggccggTAATACGACTCACTATAGGGATGCgcgaagctcgtctgtactatggccNNNNNNNNNNNNNNNNNNNNATGCccggccgcttcgagcagacatgata |
| GB_34 | | tatcatgtctgctcgaagcggccggGCAT |

**Table S6 – PCR Primer sequences**

| | Cell type | slope | intercept |
|---|---|---|---|
| **R1 to R2** | HepG2 | 0.626018 | -1.524307 |
| | K562 | 0.614353 | -1.288952 |
| **R0-MPRA to R1** | HepG2 | 0.897912 | -0.691838 |
| | K562 | 0.883498 | -1.052554 |

**Table S7 – Batch correction regression coefficients**
